# Deep Plasma Proteome Profiling by Modulating Single Nanoparticle Protein Corona with Small Molecules

**DOI:** 10.1101/2024.03.06.582595

**Authors:** Ali Akbar Ashkarran, Hassan Gharibi, Seyed Amirhossein Sadeghi, Seyed Majed Modaresi, Qianyi Wang, Teng-Jui Lin, Ghafar Yerima, Ali Tamadon, Maryam Sayadi, Maryam Jafari, Zijin Lin, Danilo Ritz, David Kakhniashvili, Avirup Guha, Mohammad R.K. Mofrad, Liangliang Sun, Markita P. Landry, Amir Ata Saei, Morteza Mahmoudi

## Abstract

The protein corona, a dynamic biomolecular layer that forms on nanoparticle (NP) surfaces upon exposure to biological fluids is emerging as a valuable diagnostic tool for improving plasma proteome coverage analyzed by liquid chromatography-mass spectrometry (LC-MS/MS). Here, we show that spiking small molecules, including metabolites, lipids, vitamins, and nutrients (namely, glucose, triglyceride, diglycerol, phosphatidylcholine, phosphatidylethanolamine, L-α-phosphatidylinositol, inosine 5′-monophosphate, and B complex), into plasma can induce diverse protein corona patterns on otherwise identical NPs, significantly enhancing the depth of plasma proteome profiling. The protein coronas on polystyrene NPs when exposed to plasma treated with an array of small molecules (n=10) allowed for detection of 1793 proteins marking an 8.25-fold increase in the number of quantified proteins compared to plasma alone (218 proteins) and a 2.63-fold increase relative to the untreated protein corona (681 proteins). Furthermore, we discovered that adding 1000 µg/ml phosphatidylcholine could singularly enable the detection of 897 proteins. At this specific concentration, phosphatidylcholine selectively depleted the four most abundant plasma proteins, including albumin, thus reducing the dynamic range of plasma proteome and enabling the detection of proteins with lower abundance. By employing an optimized data-independent acquisition (DIA) approach, the inclusion of phosphatidylcholine led to the detection of 1436 proteins in a single plasma sample. Our molecular dynamic results revealed that phosphatidylcholine interacts with albumin via hydrophobic interactions, h-bonds, and water-bridges. Addition of phosphatidylcholine also enabled the detection of 337 additional proteoforms compared to untreated protein corona using a top-down proteomics approach. These significant achievements are made utilizing only a single NP type and one small molecule to analyze a single plasma sample, setting a new standard in plasma proteome profiling. Given the critical role of plasma proteomics in biomarker discovery and disease monitoring, we anticipate widespread adoption of this methodology for identification and clinical translation of proteomic biomarkers into FDA approved diagnostics.

## Introduction

The quest to comprehensively analyze the plasma proteome has become crucial for advancing disease diagnosis and monitoring, as well as biomarker discovery.^1, 2^ Yet, obstacles like identifying low-abundance proteins remain owing to the prevalence of high-abundance proteins in plasma where the seven most abundant proteins collectively represent 85% of the total protein mass.^3, 4^ Peptides from these high-abundance proteins, especially those of albumin, tend to dominate mass spectra impeding the detection of proteins with lower-abundance.

To address this challenge, techniques such as affinity depletion, protein equalizer, and electrolyte fractionation have been developed to reduce the concentration of these abundant proteins, thereby facilitating the detection of proteins with lower-abundance.^5, 6, 7^ Additionally, a range of techniques has been developed to enhance the throughput and depth of protein detection and identification, from advanced acquisition modes to methods that concentrate low-abundance proteins or peptides for liquid chromatography mass spectrometry (LC-MS/MS) analysis.^5, 8, 9, 10, 11, 12, 13^ For instance, in the affinity depletion strategy^14^, affinity chromatography columns are used with specific ligands that bind to high-abundance proteins such as albumin, immunoglobulins, and haptoglobin. However, cost and labor associated with such depletion strategies hampers their application for large cohorts. As another example, salting-out technique^15^ is used to add reagents (e.g., ammonium sulfate) to selectively precipitate high-abundance proteins, leaving the lower-abundance proteins in the supernatant. However, these methods can introduce biases in precipitating lower-abundance proteins as well, therefore, additional robust strategies are needed to ensure low-abundance proteins with high diagnostic potential are not missed in biomarker discovery studies. More details on the limitations of these strategies are presented elsewhere.^16^

Recently, nanoparticles (NPs) have gained attention for their ability to support biomarker discovery through analysis of the spontaneously-forming protein/biomolecular corona (i.e., a layer of biomolecules, primarily proteins, that forms on NPs when exposed to plasma or other biological fluids)^5, 17, 18, 19, 20, 21, 22, 23, 24, 25^. The protein corona can contain a unique ability to concentrate proteins with lower abundance, easily reducing the proteome complexity for LC-MS/MS analysis.^5, 17, 26^ While the physicochemical properties of NPs do indeed influence the structure of their protein corona, it is generally observed that nanoscale materials exhibit different protein abundances compared to the original plasma protein composition.^27^ In essence, most NPs have the potential to form a protein corona with a distinct protein composition and abundance, differing from the native plasma proteins.^27^

Application of single NPs for biomarker discovery has limitations in achieving deep proteome coverage, typically enabling the detection of only hundreds of proteins.^28^ To enhance proteome coverage and quantify a higher number of plasma proteins, the use of a protein corona sensor array or multiple NPs with distinct physicochemical properties can be implemented. This approach leverages the unique protein corona that forms on each NP to increase proteome coverage, but carries the drawback of having to analyze multiple NP samples and needing to test many NP types to reach the desired depth.^5, 26, 29^ In addition, the use of single NPs offers several advantages over multiple NPs, particularly in terms of commercialization and the regulatory complexities associated with multi-NP systems.^30^ Additionally, utilizing a single type of NP can streamline the MS analysis process, reducing the time required to analyze large cohorts in plasma proteomics studies.

Small molecules native to human biofluids play a significant role in regulating human physiology, often through interactions with proteins. Therefore, we hypothesize that small molecules might influence the formation of the NP protein corona and serve to enrich specific proteins including biomarkers or low abundance proteins. Recent findings have reported that high levels of cholesterol results in a protein corona with enriched apolipoproteins and reduced complement proteins, which is due to the changes in the binding affinity of the proteins to the NPs in the presence of cholesterol.^31^ Accordingly, we hypothesized that small molecules endogenous to human plasma may affect the composition of the NP protein corona differently depending on whether these molecules act individually or collectively.^32^

Our work presents an efficient methodology that harnesses the influence of various small molecules in creating diverse protein coronas on otherwise identical polystyrene NPs. Our primary hypothesis, corroborated by our findings, posits that introducing small molecules into plasma alters the manner in which the plasma proteins engage with NPs. This alteration, in turn, modulates the protein corona profile of the NPs. As a result, when NPs are incubated with plasma pre-treated with an array of small molecules at diverse concentrations, these small molecules significantly enhance the detection of a broad spectrum of low-abundance proteins through LC-MS/MS analyses. The selected small molecules include essential biological metabolites, lipids, vitamins, and nutrients consisting of glucose, triglyceride, diglycerol, phosphatidylcholine (PtdChos), phosphatidylethanolamine (PE), L-α-phosphatidylinositol (PtdIns), inosine 5′-monophosphate (IMP), and B complex and their combinations. The selection of these molecules was based on their ability to interact with a broad spectrum of proteins, which significantly influences the composition of the protein corona surrounding NPs. For example B complex components can interact with a wide range of proteins including albumin^33, 34^, hemoglobin^33^, myoglobin^35^, pantothenate permease^36^, acyl carrier protein^37^, lactoferrin^38^, prion^39^, β-amyloid precursor^40^, and niacin-responsive repressor^41^. Additionally, to assess the potential collective effects of these molecules, we analyzed two representative "molecular sauces." Molecular sauce 1 contained a blend of glucose, triglyceride, diglycerol, PtdChos, and molecular sauce 2 consisted of PE, PtdIns, IMP, and vitamin B complex.

Why did we choose polystyrene NPs for this study? Our team has extensive experience in analyzing the composition and profiles of the protein corona on various types of NPs, including gold^42, 43, 44^, superparamagnetic iron oxide^45, 46, 47^, graphene oxide^48, 49, 50^, iron-platinum ^51^, zeolite ^52, 53^, silica^54, 55^, polystyrene^54, 56, 57, 58^, silver^59^, and lipids^22, 60, 61^. In this study, we specifically selected highly uniform polystyrene NPs for two primary reasons: i) polystyrene NPs have a protein corona that encompasses a broad spectrum of protein categories, including immunoglobulins, lipoproteins, tissue leakage proteins, acute phase proteins, complement proteins, and coagulation factors. This diversity is crucial for achieving wide proteome identification, which is essential for our research objectives; ii) these particles are tested widely for numerous applications in nanobiomedicine: we^56, 57, 58^ and other groups^62, 63, 64, 65, 66^ have conducted extensive optimization, employing a wide range of characterizations, including MS, to analyze the protein corona of polystyrene NPs. This rigorous optimization ensures highly accurate and reproducible results.

Our findings confirm that the addition of these small molecules in plasma generates distinct protein corona profiles on otherwise identical NPs, significantly expanding the range of the plasma proteome that can be captured and detected by simple LC-MS/MS analysis. Notably, we discover that the addition of specific small molecules, such as PtdChos, leads to a substantial increase in proteome coverage, which is attributed to the unique ability of PdtChos to bind albumin and reduce its participation in protein corona formation. Therefore, PtdChos coupled with NP protein corona analysis can replace the expensive albumin depletion kits and accelerate the plasma analysis workflow by reducing processing steps. Furthermore, our single small molecule-single NP platform reduces the necessity for employing multiple NP workflows in plasma proteome profiling. This approach can seamlessly integrate with existing LC-MS/MS workflows to further enhance the depth of plasma proteome analysis for biomarker discovery.

## Results

### Protein corona and small molecules enable deep profiling of the plasma proteome

We assessed the effect of eight distinct small molecules, namely, glucose, triglyceride, diglycerol, PtdChos, PE, PtdIns, IMP, and vitamin B complex, on the protein corona formed around polystyrene NPs. The workflow of the study is outlined in **Supplementary Scheme 1**.

Commercially available plain polystyrene NPs, averaging 80 nm in size, were purchased. Each small molecule, at varying concentrations (10, 100, and 1000 µg/ml; we selected a broad range of small molecule concentrations to determine the optimal levels for maximizing proteome coverage), was first incubated with commercial pooled healthy human plasma at 37°C for 1 h allowing the small molecules to interact with the biological matrix. The concentration of each small molecule was carefully adjusted to ensure that the final concentration in the combined molecular solutions were 10, 100, or 1000 µg/ml for each component, consistent with the concentration used for individual small molecules. Subsequently, NPs at a concentration of 0.2 mg/ml were introduced into the plasma containing small molecules or sauces and incubated for an additional hour at 37°C with agitation. It is noteworthy that the NPs concentration was chosen in a way to avoid any protein contamination (which was detected at concentrations of 0.5 mg/ml and higher) in the protein corona composition, which may cause errors in the proteomics data.^67, 68^ These methodological parameters were refined from previous studies to guarantee the formation of a distinct protein corona around the NPs. **Supplementary Fig. 1** offers further details on our methodologies, showcasing dynamic light scattering (DLS), zeta potential, and transmission electron microscopy (TEM) analyses for both the untreated NPs and those covered by a protein corona^69^. The untreated polystyrene NPs exhibited excellent monodispersity, with an average size of 78.8 nm with the polydispersity index of 0.026 and a surface charge of -30.1 ± 0.6 mV. Upon formation of the protein corona, the average size of NPs expanded to 113 nm, and the surface charge shifted to -10 mV ± 0.4 mV. TEM analysis further corroborated the size and morphology alterations of the NPs before and after protein corona formation (**Supplementary Fig. 1**).

To investigate how spiking different concentrations of small molecules can influence the molecular composition of the protein corona, samples were subjected to LC-MS/MS analysis for high-resolution proteomic analysis. While the analysis of plasma alone led to quantification of 218 unique proteins, analysis of the protein corona formed on the polystyrene NPs significantly enhanced the depth of plasma proteome sampling to enable quantification of 681 unique proteins. Furthermore, the inclusion of small molecules further deepened plasma proteome sampling to enable quantification of between 397 and up to 897 unique proteins, depending on the small molecules added to plasma prior to corona formation. When comparing the use of protein coronas, both with and without the inclusion of small molecules, to the analysis of plasma alone (**Fig. 1a** and **Supplementary Data 1**), there is a notable increase— approximately a threefold rise—in the number of proteins that can be quantified. The CVs of the number of quantified proteins between 3 technical replicates were generally less than 1.54% for all sample types (**Supplementary Table 1**).

**Fig. 1.**
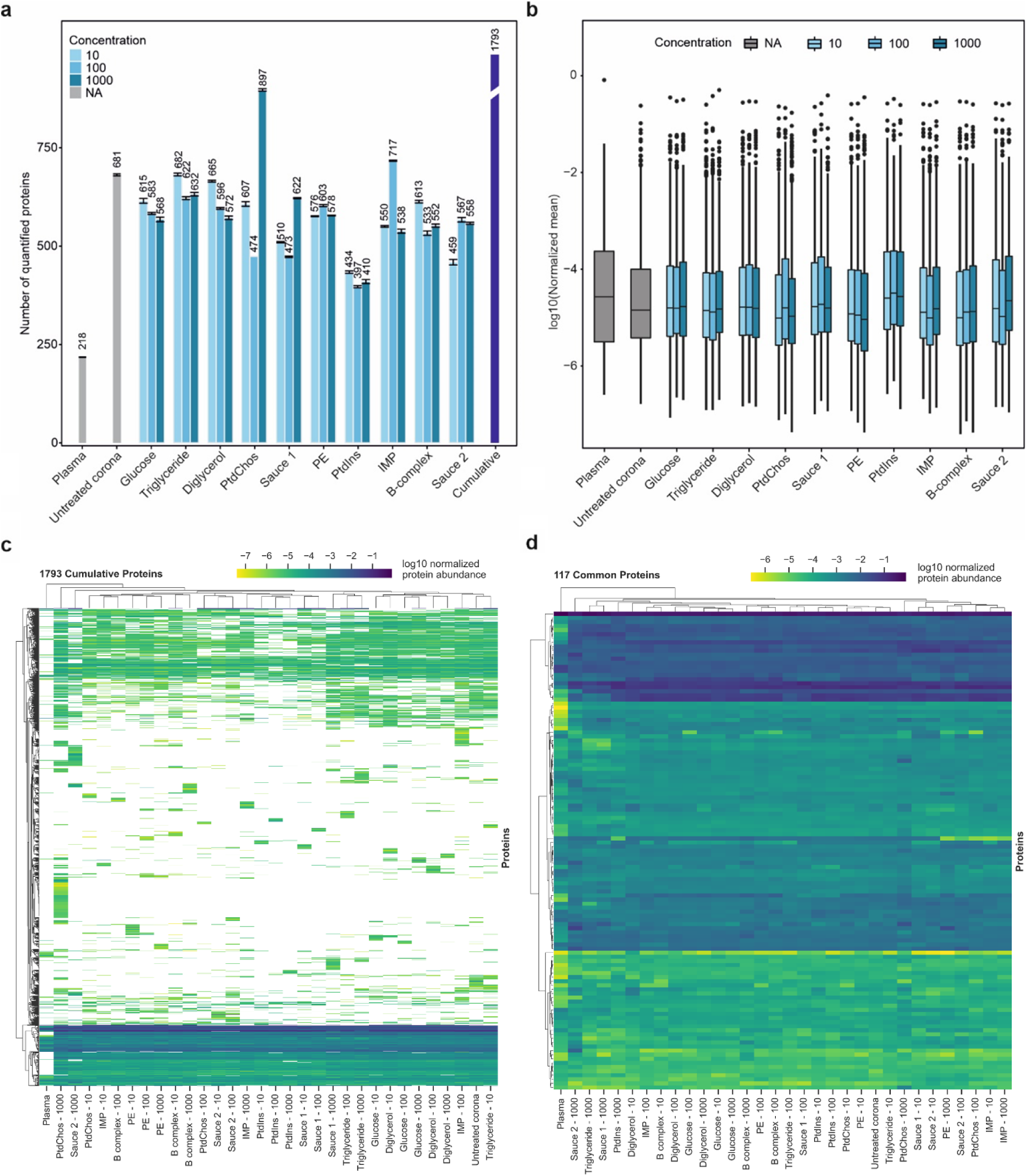
Small molecules affect the plasma proteome sampling. **a,** The number of quantified proteins in plasma, untreated protein corona and protein coronas in the presence of small molecules and molecular sauces (mean ± SD of three technical replicates). The cumulative number of unique proteins identified using untreated protein corona and corona treated with various small molecules is also shown using the purple bar. For fair comparison, the database was performed individually for each small molecule. **b,** The distribution of averaged normalized abundances of three technical replicates for proteins quantified in the plasma, untreated protein corona and protein coronas in the presence of small molecules and molecular sauces (center line, median; box limits contain 50%; upper and lower quartiles, 75 and 25%; maximum, greatest value excluding outliers; minimum, least value excluding outliers; outliers, more than 1.5 times of upper and lower quartiles). **c,** Clustered heatmap of normalized abundance of all 1793 proteins quantified across all samples. White denotes not detected. **d,** Clustered heatmap of normalized abundance of 117 shared proteins across all samples. Experiments were performed in three technical replicates.

Interestingly, the concentration of small molecules did not significantly affect the number of quantified proteins in a concentration-dependent manner; only a small stepwise reduction in the number of quantified proteins was noted with increasing concentrations of glucose and diglycerol. Cumulatively, the incorporation of small molecules and molecular sauces into the protein corona of NPs led to a significant increase in protein quantification, with a total of 1793 proteins identified, marking an 8.25-fold increase compared to plasma proteins alone. Specifically, the addition of small molecules resulted in the quantification of 1573 additional proteins compared to plasma alone, and 1037 more proteins than the untreated protein corona. Strikingly, spiking 1000 µg/ml of PtdChos increased the number of quantified proteins to 897 (1.3-fold of quantified proteins in untreated plasma), singlehandedly. This observation prompted a detailed investigation into the influence of PtdChos on plasma proteome coverage, which is elaborated in the following sections. It noteworthy that the superior performance of PtdChos alone compared to Molecular Sauce 1 could be attributed to interactions between the small molecules in the mixture, which may have lowered the effective concentration of PtdChos (for example, the interactions between PtdChos and triglycerides)^70, 71^. Mass spectrometry workflow and the type of data analysis have a critical influence on proteomics outcomes in general^9, 72, 73, 74, 75^, as well as in the specific field of protein corona research^57, 58, 76^. For instance, our recent study demonstrated that identical corona-coated polystyrene NPs analyzed by different mass spectrometry centers resulted in a wide range of quantified proteins, varying from 235 to 1,430 (5.1 fold increase as compared to plasma alone).^57^ To mitigate the impact of these variables on the interpretation of how small molecules can enhance proteome coverage, we chose to report our data as fold changes of the number of quantified proteins relative to control plasma and untreated corona samples. This approach offers a more objective assessment of the role of small molecules in enhancing proteome analysis, minimizing the confounding effects of different workflows and data analysis techniques that may be employed by various researchers.

The distribution of normalized protein intensities for the samples is shown in **Fig. 1b**. The median value in the plasma group was notably higher than in the other samples, although the overall distribution did not differ significantly. In general, the proteomes obtained from protein corona profiles in the presence of small molecules showed a good correlation (generally a Pearson correlation above 0.6 for most small molecule comparisons) demonstrating the faithful relative representation of proteins after treatment with different small molecules (**Supplementary Fig. 2**).

### Small molecules diversify the protein corona composition

We next investigated if the addition of small molecules would change the type and number of proteins detected by LC-MS/MS. Indeed, each small molecule and the molecular sauces generated a proteomic fingerprint that was distinct from untreated protein corona or those of other small molecules (**Fig. 1c**). Spiking small molecules led to detection of a diverse set of proteins in the plasma. Interestingly, even different concentrations of the same small molecules or molecular sauces produced unique fingerprints. A similar analysis was performed for the 117 shared proteins across the samples (**Fig. 1d**). The Venn diagrams in **Supplementary Fig. 3a** and **3b** show the number of unique proteins that were quantified in the respective group across all concentrations which were not quantified in the plasma or in the untreated protein corona. These results suggest that spiking small molecules into human biofluids can diversify the range of proteins that are identifiable in protein corona profiles, effectively increasing proteomic coverage to lower abundance proteins. Such an enrichment or depletion of a specific subset of proteins can be instrumental in biomarker discovery focused on a disease area. This feature can also be used for designing assays where the enrichement of a known biomarker is facilitated by using a given small molecule. As representative examples, a comparison of enriched and depleted proteins for molecular sauce 1 and 2 against the untreated protein corona is shown in **Supplementary Fig. 3c** and **3d**, respectively (**Supplementary Data 2**). In certain cases, the enrichment or depletion was drastic, spanning several orders of magnitude. The enriched and depleted proteins for the molecular sauce 1 and 2 were mapped to KEGG pathways and biological processes in StringDB (**Supplementary Fig. 3c-d**). While most of the enriched pathways were shared, some pathways were specifically enriched for a given molecular sauce. For example, systemic lupus erythematosus (SLE) was only enriched among the top pathways for molecular sauce 2. Therefore, the small molecules can be potentially used for facilitating the discovery of biomarkers for specific diseases, or for assaying the abundance of a known biomarker in disease detection.

Similar analyses were performed for all the small molecules and the volcano plots for the highest concentration of each molecule (i.e., 1000 µg/ml) are demonstrated in **Supplementary Fig. 4** (**Supplementary Data 2**). A pathway analysis was also performed for all the significantly changing proteins for each small molecule at all concentrations (**Supplementary Fig. 5**). To facilitate comparison, we have combined the enrichement analysis for all the samples vs. the untreated protein corona in **Supplementary Fig. 6**.

To demonstrate how small molecules affect the composition and functional categories of proteins in the protein corona, potentially aiding in early diagnosis of diseases (since proteins enriched in the corona are pivotal in conditions like cardiovascular and neurodegenerative diseases), we utilized bioanalytical methods^77^ to categorize the identified proteins based on their blood-related functions namely complement activation, immune response, coagulation, acute phase response, and lipid metabolism (**Supplementary Fig. 7**). In our analysis, apolipoproteins were major protein types that were found in the small molecule treated protein corona, and their types and abundance were heavily dependent to the type and concentrations of the employed small molecules (**Supplementary Fig. 8**). Similarly, the enrichment of other specific protein categories on NPs surfaces was influenced by the type and concentration of small molecules used (**Supplementary Fig. 8**). For example, antithrombin-III in coagulation factors plays a significant role in the protein corona composition of all tested small molecules, but this effect is observed only at their highest concentration. At lower concentrations, or in the untreated protein corona, this considerable participation is not evident (**Supplementary Fig. 8**). This ability of small molecules to modify the protein composition on NPs highlights their potential for early disease diagnosis (e.g., apolipoprtoeins in cardiovascular and neurodegenerative disorders)^78, 79^, where these protein categories are crucial in disease onset and progression ^78^.

### PtdChos reduces the plasma proteome dynamic range and increases proteome coverage by depleting the abundant plasma proteins

To understand whether the quantification of a higher number of proteins in protein corona profiles was due to a lower dynamic range of proteins available in human plasma for NP binding, we plotted the maximum protein abundance vs. minimum protein abundance for plasma alone, and plasma treated with small molecules in **Supplementary Fig. 9**. The plasma alone showed the highest dynamic range, suggesting that identification of low-abundance proteins would be most difficult from plasma alone. Conversely, addition of small molecules was shown to reduce plasma protein dynamic range, thereby allowing for detection of more peptides and quantification of proteins with lower abundance through the NP protein corona.

Notably, while albumin accounted for over 81% of our plasma sample, its representation was significantly lowered to an average of 29% in the protein coronas, both with and without small molecule modifications. This reduction was most pronounced with PtdChos treatment at 1000 µg/ml, where albumin levels dropped to around 17% of plasma proteins (**Fig. 2a**). Despite these changes, albumin remained the most abundant protein in all samples. A similar diminishing trend was observed for the second and third most abundant proteins, serotransferrin (TF) and haptoglobin (HB), which make up about 3.9% and 3.6% of plasma protein abundance, respectively. The rankings of these proteins’ abundance in each sample are depicted above the panels in **Fig. 2a**. From this analysis, it is evident that the protein corona, both in its native form and when altered by small molecules, can drastically reduce the combined representation of the top three proteins from about 90% to roughly 29%. The most substantial reduction was observed with PtdChos at 1000 µg/ml, reducing the top three proteins’ cumulative representation from 90% to under 17%. PtdChos treatment also effectively reduced the levels of the fourth most abundant plasma protein IGHA1. This significant decrease in the abundance of highly prevalent plasma proteins explains the marked increase in the number of unique proteins detected from NP corona samples treated with PtdChos (897 proteins identified in the PtdChos-treated protein corona vs. 681 proteins identified in the untreated corona vs. 218 proteins identified in the untreated plasma, as shown in **Fig. 1a**). These results indicate that high concentrations of PtdChos can be strategically employed to enable more comprehensive plasma protein sampling by specifically targeting and depleting the most abundant plasma proteins, especially albumin.

**Fig. 2.**
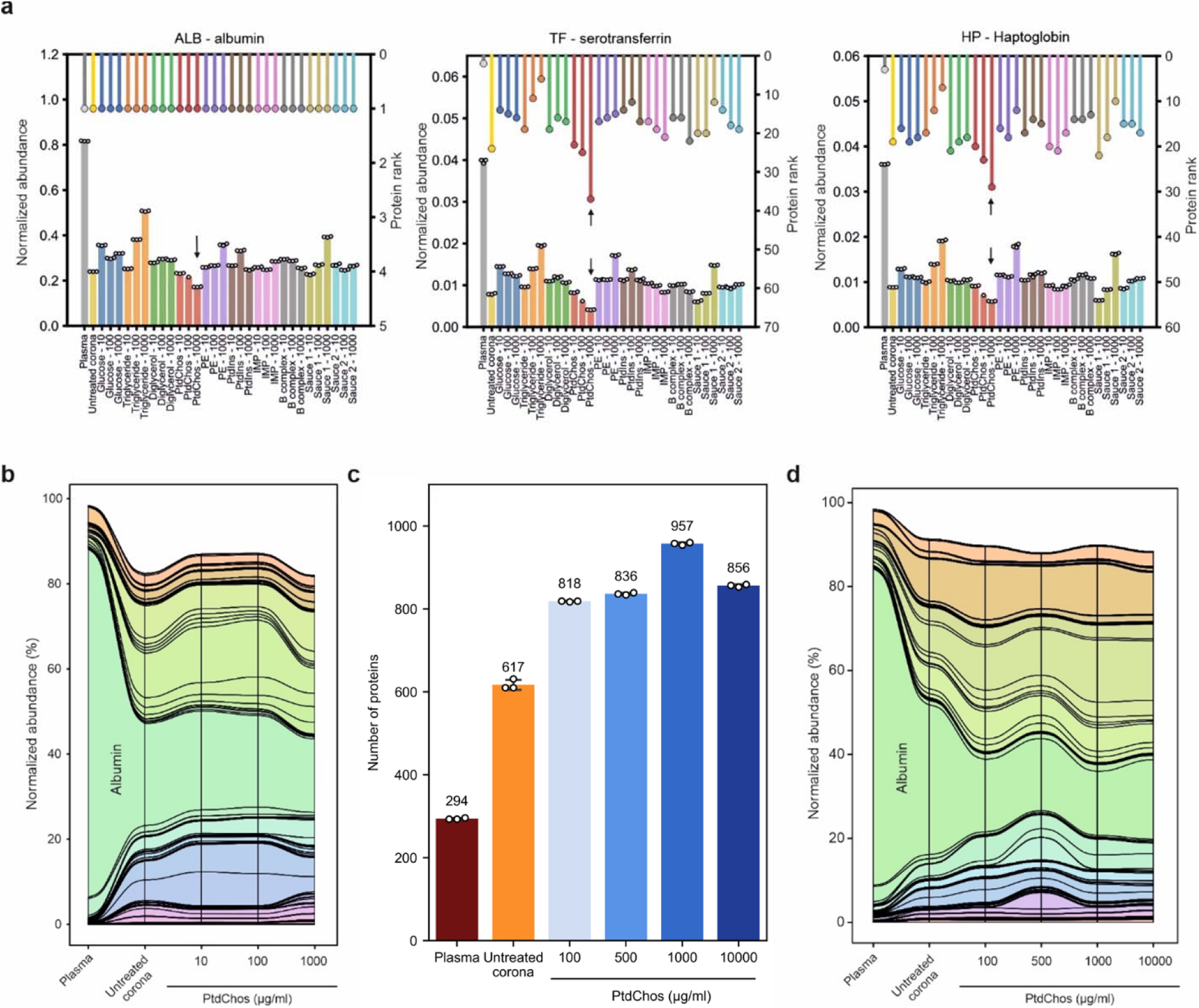
PtdChos can deplete the most abundant plasma proteins in protein corona profiles. **a,** Normalized protein abundance (left axis, bar plot) and protein rankings (right axis, lollipop plot) in untreated plasma, untreated protein corona, and small-molecule treated protein corona. **b**, A stream (or alluvial) diagram illustrating the significant depletion of abundant plasma proteins, particularly albumin, following the incubation of plasma with NPs and PtdChos (only shared proteins with plasma are included). **c**, Total count of proteins identified in plasma, untreated protein corona, and protein corona treated with PtdChos at various concentrations (mean ± SD of three technical replicates). **d**, A stream diagram demonstrating the depletion pattern of abundant plasma proteins, especially albumin, in response to NP addition and enhanced with escalating concentrations of PtdChos (only shared proteins with plasma are included).

The stream (or alluvial) diagram in **Fig 2b** shows the overall changes in the representation of proteins found in plasma upon incubation of protein corona with different concentrations of PtdChos. To validate this discovery, we prepared fresh samples treated with a series of PtdChos concentrations ranging from 100 µg/ml to 10,000 µg/ml (**Supplementary Data 3**). As shown in **Fig 2c**, 957 proteins could be quantified in the protein corona treated with PtdChos at 1000 µg/ml, while neither lower concentration nor further addition of PtdChos did not enhance the number of quantified proteins. The CVs of the number of quantified proteins between 3 technical replicates were generally less than 2% for all sample types (**Supplementary Table 2**). The stream diagram in **Fig 2d** shows the specific depletion of albumin and a number of other abundant proteins in plasma upon addition of PtdChos, allowing for more robust detection of other proteins with lower abundance.

To confirm that the improved proteome coverage achieved with PtdChos treatment is independent of the LC-MS platform or the data acquisition mode used, we prepared new samples of plasma, untreated protein corona, and protein corona treated with 1000 µg/ml PtdChos, and analyzed them using LC-MS in the DIA mode. We identified 322 proteins in the plasma alone, 1011 proteins in the untreated protein corona samples, and 1436 proteins in the protein corona treated with PtdChos (1.4-fold increase over the untreated corona) (**Supplementary Data 4**). These findings not only validate the enhancement of plasma proteome coverage by PtdChos but also illustrate the capability of PtdChos to facilitate the in-depth profiling of the plasma proteome associated with protein corona formed on the surface of a single type of NP. Since the ratio of the number of quantified proteins through PtdChos spiking is generally around 1.4-fold higher than in the NP corona alone, PtdChos can be incorporated into any LC-MS workflow aiming to boost plasma proteome profiling. More optimized plasma proteomics pipelines, TMT multiplexing coupled to fractionation, or high-end mass spectrometers such as Orbitrap Astral are envisioned to quantify an even higher number of proteins that those reported in the current study.

To confirm the role of PtdChos in enhancing the proteome depth of the protein corona, we expanded our analysis by using additional NPs and four plasma samples from individual donors. Specifically, we tested seven additional commercially available and highly uniform NPs with distinct physicochemical properties: polystyrene NPs of varying sizes (mean diameters of 50, 100, and 200 nm) and surface charges (carboxylated and aminated polystyrene NPs, both with the mean diameter of 100 nm), as well as silica NPs with the mean sizes of 50 and 100 nm. These NPs have been extensively characterized and widely utilized for protein corona analysis by numerous research groups including our own.^54, 80, 81, 82, 83, 84, 85^

The protein corona samples from different NPs were analyzed in the DIA mode with the 30 samples per day (SPD) setting with 44 min acquisition time. Our analysis revealed two key findings: (i) the physicochemical properties of NPs significantly influence the effectiveness of PtdChos in enhancing the number of quantified proteins in plasma, and (ii) incorporating additional plasma samples can markedly increase the overall number of identified proteins (**Supplementary Data 5**; **Supplementary Fig. 10a-b**). Polystyrene NPs, in general and due to their hydrophobic nature, showed higher protein detection capacity than silica NPs (*p* value = 0.012; Student’s t-test, two-sided with unequal variance). The average number of quantified proteins using polystyrene NPs was 823.4 vs. 633 with silica NPs, while cumulatively there were 1241 unique quantified proteins in polystyrene NPs compared to 1024 in silica. Polystyrene NPs with 200 nm size provided the highest proteome coverage, although the difference in the number of quantified proteins was comparable to the same type of NPs with other sizes. Plain and positively charged polystyrene NPs had a better performance than carboxylated NPs. Our analysis also revealed the inter-individual variabilities between patients. The percentage CVs of the number of proteins quantified across four donors were generally lower for polystyrene NPs than silica NPs (14.4 vs. 21.6%) (**Supplementary Table 3**).

### PtdChos increases the number of detected plasma proteoforms

We then asked if PtdChos can enhance the number of detected proteoforms in top-down proteomics as well. Proteoforms represent distinct structural variants of a protein product from a single gene, including variations in amino acid sequences and post-translational modifications.^86^ Proteoforms originating from the same gene can exhibit divergent biological functions and are crucial for modulating disease progression.^87, 88, 89^ Therefore, proteoform-specific measurement of the protein corona, along with their improved detection depth through the use of small molecules, will undoubtedly provide a more accurate characterization of the protein molecules within the corona. We compared the chromatogram, the number of proteoform identifications, proteoform mass distribution, and differentially represented proteins between the untreated corona and PtdChos-treated samples. The LC-MS/MS data showed consistent base peak chromatograms, the number of proteoform identifications, and the number of proteoform-spectrum matches (PrSMs) across the technical triplicates of both the control and PtdChos-treated samples (**Fig. 3a** and **b**, respectively). However, the treated sample exhibited a significant signal corresponding to the small molecule after 60 min of separation time (**Fig. 3b**), validating our hypothesis that small molecules interact with plasma proteins, causing the observed variation in the protein corona on the NPs’ surface. In total, 637 proteoforms were identified across the two samples (with technical triplicates for each sample) (**Fig. 3c**). Data analysis using Perseus software (Version 2.0.10.0) revealed that only 110 proteoforms overlapped between the two samples (the minimum number of valid values for filtering data was set to 1).

**Fig. 3.**
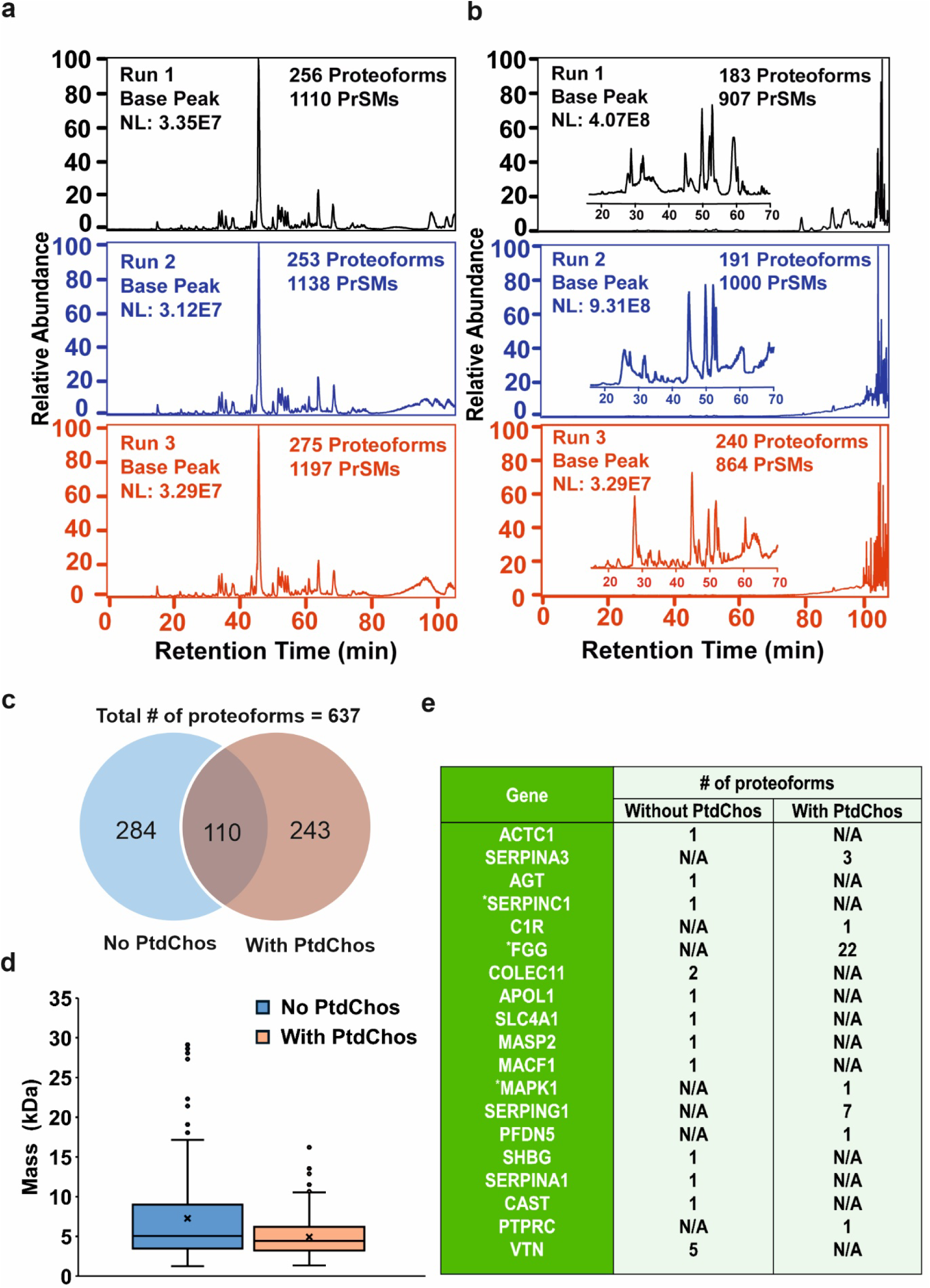
Base peak chromatograms and proteoform analysis of protein corona samples. **a,** Base peak chromatogram of eluted protein corona without PtdChos and **b,** with PtdChos in healthy human plasma after RPLC-MS/MS analyses. Two protein corona samples were prepared in parallel and analyzed by RPLC-MS/MS, with each sample measured in triplicate. **c,** The number of proteoform identifications in each sample and the overlap of proteoform identifications between the two samples. **d,** Mass distribution of proteoforms between the two samples, with the cross sign representing the mean proteoform mass in each sample (Center line—median; box limits contain 50% of data; upper and lower quartiles, 75 and 25%; maximum—greatest value excluding outliers; minimum—least value excluding outliers; outliers— more than 1.5 times of the upper and lower quartiles). **e,** Summary of some disease-related protein biomarkers identified by top-down proteomics. The Genes were determined according to the information in the Human Protein Atlas (https://www.proteinatlas.org/) and three genes labelled by * represent FDA approved drug target.

The proteoform mass distribution differed between the two samples (**Fig. 3d**). Although the average proteoform masses were similar, the box plot indicated a greater number of larger proteoform identifications in the control sample (over 20 kDa). We hypothesize that PtdChos can bind to large proteins, and due to the high concentration of PtdChos relative to the proteoforms, the signals of these large proteoforms may be obscured.

Additional data analyses identified differential proteins in this study (**Fig. 3e**). The top-down proteomics approach identified specific gene products that bind to the NP surface in the presence of PtdChos.

### PtdChos interacts with human serum albumin via hydrophobic interactions, h-bonds, and water-bridges

To determine the types of interactions between albumin and PtdChos, we conducted all-atoms molecular dynamics (MD) simulations with various numbers of PtdChos molecules (**Supplementary Fig. 11**). First, we performed blind and site-specific molecular docking simulations to find the most favorable binding sites for PtdChos on albumin. We then used the top 10 most favorable non-overlapping binding poses, as quantified by binding affinity, for our MD simulations (**Supplementary Fig. 11b-c**). Four types of systems with the top 1, 3, 5, and 10 PtdChos molecules, respectively, were investigated via 100 ns simulations. As evidenced by the sum of Lennard-Jones and coulombic interaction energies shown on **Fig. 4a**, PtdChos strongly interacts with albumin. A nearly additive effect occurs from 1 to 3 ligands added. However, the 5 ligands system has a similar total energy as the 3 ligands one. This may indicate that some PtdChos molecules do not strongly interact with albumin. When the number of ligands increased to 10, we noticed an almost 2-fold increase in energy as compared to the 5 ligands system. To further quantify the strength of interactions between albumin and PtdChos, we calculated the effective free energy of the 4 types of systems, obtaining a similar trend (**Fig. 4b**). The average root mean square fluctuations of albumin residues reveals consistent peaks with increase in fluctuations as the number of ligands increases (**Fig. 4c**). This may suggest that the protein conformation does not change drastically based on the number of ligands added. The average root mean square deviations of the PtdChos heavy atoms show similar values for the 1 and 3 ligand systems but higher values for 5 and 10 ligands systems (**Fig. 4d**). This confirms that the first few poses form a more stable interaction with albumin. Finally, **Fig. 4e** shows that albumin and PtdChos interact primarily via hydrophobic interactions, hydrogen bonding, and water-bridges. The hydrophobic interactions formed between albumin and the long fatty acid chains are present throughout every simulation. On the other hand, the number of hydrogen bonds and water-bridges increase significantly from the 1 ligand systems to the 3, 5, and 10 ligands systems. These interactions are mainly due to the phosphate group oxygen atoms.

**Fig. 4.**
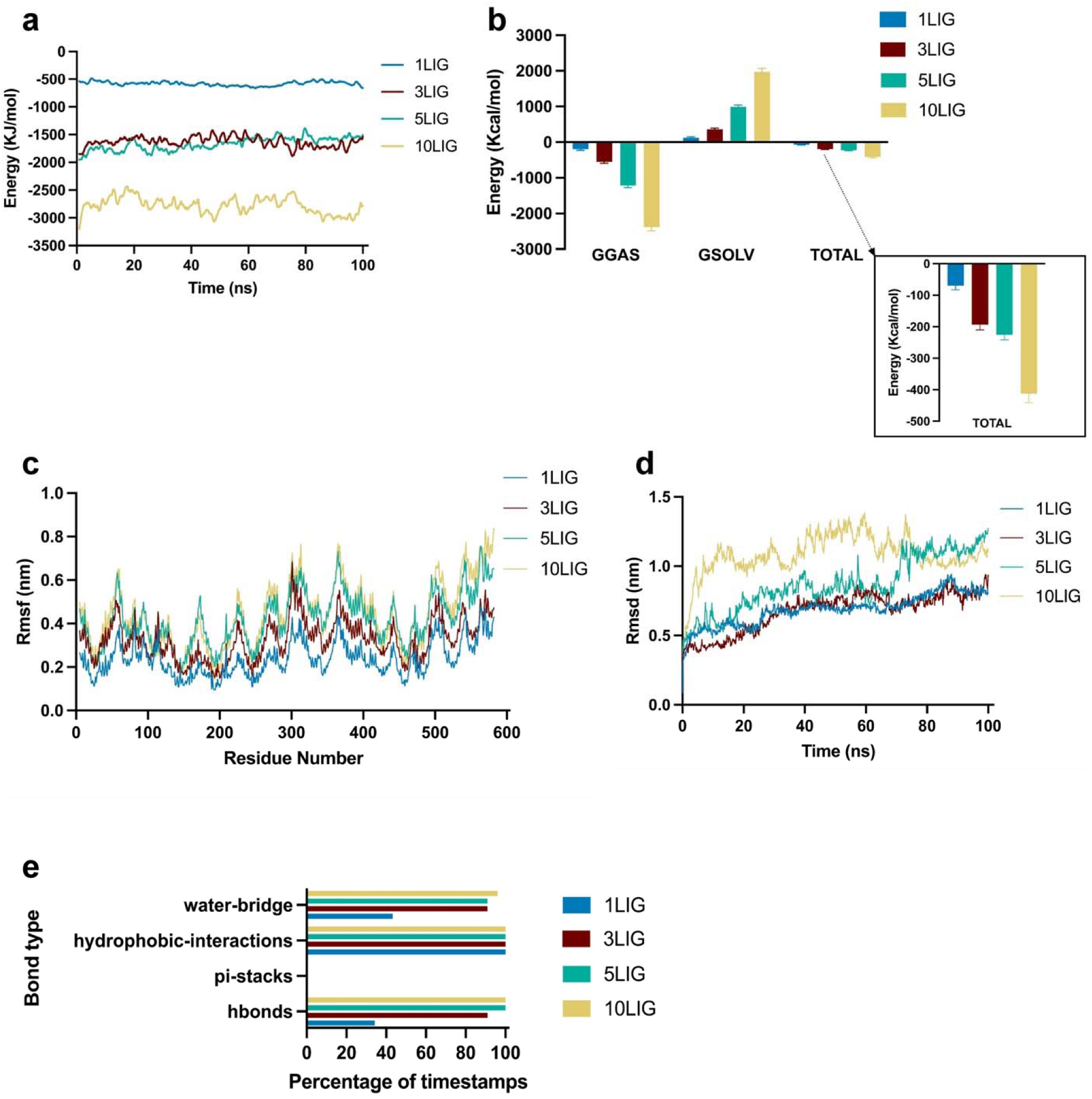
PtdChos and albumin interaction analysis in all-atoms molecular dynamics simulations. **a,** Total linear interaction energies between albumin and various number of ligands systems over simulation time. The total energy represents the sum of Lennard-Jones and coulombic energies. **b,** Effective free energy components for the different systems. GGAS represents the energy of the gas phase, GSOLV, the energy of solvation, and TOTAL is the sum of the two. Error bars represent mean ±SD. **c,** Average root mean square fluctuation of albumin residues for the 4 systems. **d,** Root mean square deviation of PtdChos over time for the 4 types of systems. **e,** Bond types that are present within each simulation. The y-axis represents the bond types, and the x-axis represents the percentage of simulation timestamps when each type of bond is present.

## Discussion

The protein corona is a layer of proteins that spontaneously adsorbs on the surface of nanomaterials when exposed to biological fluids.^90^ The composition and dynamic evolution of the protein corona is critically important as it can impact the interactions of NPs with biological systems (e.g., activate the immune system), can cause either positive or adverse biocompatibility outcomes, and can greatly affect NP biodistribution *in vivo*.^90^ The specific proteins that adsorb on the surface of the NPs depend on various factors, including the physicochemical properties of the NPs and the composition of the surrounding biological fluid.^57^ Metabolites, lipids, vitamins, nutrition, and other types of small biomolecules present in the biological fluid can interact with proteins in these fluids and influence their behavior, including their adsorption onto NPs. For example, it was shown that addition of glucose and cholesterol to plasma can alter the composition of protein corona on the surface of otherwise identical NPs.^31, 69^ Small molecules can alter the protein corona of NPs, after interaction with plasma proteins, due to various mechanisms such as i) their competition with proteins for binding to the surface of NPs; ii) altering proteins’ binding affinities to NPs; and iii) changing protein conformation.^31, 69^ For example, previous studies revealed that triglyceride, PtdChos, PE, and PtdIns can interact with lipovitellin^91^, C-reactive protein^92^, protein Z^93^, and myelin basic protein (MBP)^94^, respectively. Each individual small molecule and their combinations interrogates tens to hundreds of additional proteins across a broad dynamic range in an unbiased and untargeted manner. Our results also suggest that endogenous small molecule function may help guide which small molecule(s) can enrich protein biomarkers of a specific disease class. Therefore, any changes in the level of the small molecules in the body can alter the overall composition of the protein corona, leading to variations in the types and number of proteins that bind to NPs and consequently their corresponding interactions with biosystems.^31, 32^

Among the various employed small molecules, we discovered that PtdChos alone demonstrates a remarkably high ability to reduce the participation of the most highly abundant proteins in protein corona composition. PtdChos is the most common class of phospholipids in the majority of eukaryotic cell membranes.^95^ For a long time, it has been established that PtdChos can engage in specific interactions with serum albumin through hydrophobic processes ^96, 97^, forming distinct protein-lipid complexes.^96, 98^ The results of our molecular dynamic evaluations of the interactions between PtdChos and albumin were in line with the literature. As a result, we found that the simple addition of PtdChos to plasma can significantly reduce albumin adsorption for the surface of polystyrene NPs, thereby creating unique opportunities for the involvement of a broader range of proteins with lower abundance in the protein corona layer. We also observed the same effects of PtdChos on enhancing the proteome coverage using different types of NPs. Not only is PtdChos is an economical and simple alternative for conventional albumin depletion strategies, but it can also deplete several other highly abundant proteins as an added advantage. This approach reduces the necessity for employing NP arrays in plasma proteome profiling, and the cost and biases that can occur with albumin depletion. Additionally, PtdChos can help accelerate plasma analysis workflows by reducing the sample preparation steps.

Our top-down proteomics analysis of both untreated and PtdChos-treated protein coronas demonstrated that incorporating small molecules such as PtdChos can significantly enhance the quantification of proteoforms within protein corona profiles. Proteoforms, which are distinct structural variants of proteins arising from genetic variations, alternative splicing, and post-translational modifications, play a crucial role in determining protein functionality and are often closely linked to disease occurrence and progression.^74, 86, 87, 99, 100^ By enriching the diversity and depth of proteoforms within the protein corona, the use of small molecules like PtdChos can substantially improve the level of information from plasma proteomics. This enhancement is particularly valuable for biomarker discovery, as the increased detection of proteoforms allows for a more nuanced understanding of disease mechanisms. The ability to capture a broader spectrum of proteoforms in the protein corona could lead to the identification of novel biomarkers that may otherwise be overlooked using traditional bottom-up proteomics approaches that mainly consider protein abundance.^74^

Our study highlights the tremendous potential of leveraging small molecules in enhancing the capabilities of protein corona profiles for broader plasma proteome analysis. By introducing individual small molecules and their combinations into plasma, we have successfully created distinct protein corona patterns on single identical NPs, thereby expanding the repertoire of attached proteins. Using our new approach, we quantified an additional 1573 unique proteins that would otherwise remain undetected in plasma. This enhanced depth in protein coverage can be attributed, in part, to the unique interactions of each small molecule, allowing for representation of a diverse set of proteins in the corona. Moreover, our findings underscore the influence of small molecules on the types and categories of proteins in the protein corona shell. This feature opens exciting possibilities for early disease diagnosis, particularly in conditions such as cardiovascular and neurodegenerative disorders, where enriched proteins, such as apolipoproteins, play pivotal roles.

However, one critical challenge that must be addressed is the standardization of proteomics analysis of the protein corona. Ensuring consistent and reproducible results across laboratories and core facilities is essential for the rapid development and successful translation of this platform into clinical applications.^57, 58, 76^ Addressing this challenge will require coordinated efforts from the scientific community to establish robust, universally accepted protocols. There are a few additional foreseeable limitations with the application of PtdChos. In certain scenarios, any depletion strategy could lead to distortion of the abundance of proteins in plasma, which can be mitigated by enforcing proper controls. Moreover, upon discovery of a biomarker, it can be validated in the cohort using orthogonal techniques such as Western blotting. Furthermore, similar to other albumin depletion strategies, certain proteins bound to albumin might be co-depleted (albuminome)^101^.

In summary, our platform is capable of quantifying up to 1793 proteins when using a single NP with an array of small molecules, while only 218 and 681 proteins could be quantified using the plasma or the NP protein corona alone. We showed the possibility of quantifying up to 1436 proteins using a single NP and PtdChos alone using a single plasma sample. Similarly in top-down proteomics, the addition of PtdChos to plasma prior to their interactions with NPs, can increase the number of quantified proteoforms in the protein corona. The cumulative number of detected proteins will therefore dramatically increase if this platform is applied to a cohort of patient samples with individual variability. Expectedly, with progressive development of both top-down and bottom-up platforms^74^, the depth of analysis can further increase towards the ultimate goal of achieving comprehensive human proteome coverage. Another alternative would be to combine our strategy with tandem mass tag (TMT) multiplexing and fractionation to achieve an even higher plasma proteome depth. We anticipate that this platform will find extensive applications in plasma proteome profiling, providing an unprecedented opportunity in disease diagnostics and monitoring.

## Materials and Methods

### Materials

Pooled healthy human plasma proteins, along with plasma from four individual healthy donors, were obtained from Innovative Research (www.innov-research.com) and diluted to a final concentration of 55% using phosphate buffer solution (PBS, 1X). Seven commercial NPs of various types (silica and polystyrene), sizes (50, 100, and 200 nm), and functional groups (plain, amino, and carboxylated) were sourced from Polysciences (www.polysciences.com). Small molecules were purchased from Sigma, Abcam, Fisher Scientific, VWR, and Beantown, and diluted to the desired concentration with 55% human plasma. Reagents for protein digestion, including guanidinium-HCl, DL-dithiothreitol (DTT), iodoacetamide (IAA), and trifluoroacetic acid (TFA), were obtained from Sigma Aldrich. Mass spectrometry-grade lysyl endopeptidase (Lys-C) was sourced from Fujifilm Wako Pure Chemical Corporation, and trypsin was obtained from Promega. Formic acid and C18 StageTips were purchased from Thermo Fisher Scientific.

### Protein corona formation on the surface of NPs in the presence of small molecules

For protein corona formation in the presence of small molecules, individual or pooled human plasma proteins 55% were first incubated with individual small molecules or in combination by preparing two molecular sauces of individual small molecules at different concentration (i.e., 10, 100, and 1000 µg/ml) for 1 h at 37 °C. Then, each type of polystyrene NPs was added to the mixture of plasma and small molecules solution so that the final concentration of the NPs was 0.2 mg/ml and incubated for another 1 h at 37 °C. It is noteworthy that all experiments are designed in a way that the concentration of NPs, human plasma, and small molecules were 0.2 mg/ml, 55%, and 10, 100, 1000 µg/ml, respectively. To remove unbound and plasma proteins only loosely attached to the surface of NPs, protein-NPs complexes were then centrifuged at 14,000’ g for 20 min, the collected NPs’ pellets were washed three times with cold PBS under the same conditions, and the final pellet was collected for further analysis.

For the PtdChos concentration study, we used various concentrations of PtdChos (i.e., 250, 750, 1000, and 10000 µg/ml) and used the same protein corona method for preparation of the samples for mass spectrometry analysis.

### NP characterization

DLS and zeta potential analyses were performed to measure the size distribution and surface charge of the NPs before and after protein corona formation using a Zetasizer nano series DLS instrument (Malvern company). A Helium Neon laser with a wavelength of 632 nm was used for size distribution measurement at room temperature. TEM was carried out using a JEM-2200FS (JEOL Ltd.) operated at 200kV. The instrument was equipped with an in-column energy filter and an Oxford X-ray energy dispersive spectroscopy (EDS) system. 20 μl of the bare NPs were deposited onto a copper grid and used for imaging. For protein corona–coated NPs, 20 μl of sample was negatively stained using 20 μl uranyl acetate 1%, washed with DI water, deposited onto a copper grid, and used for imaging. PC composition was also determined using LC-MS/MS.

### Bottom-up LC-MS/MS sample preparation for the screening and concentration series experiments

The collected protein corona-coated NP pellets were resuspended in 20 µL of PBS containing 0.5M guanidinium-HCl. The proteins were reduced with 2 mM DTT at 37°C for 45 min and then alkylated with 8 mM IAA for 45 min at room temperature in the dark. Subsequently, 5 µL of LysC at 0.02 µg/µL in PBS was added and incubated for 4 h, followed by the addition of the same concentration and volume of trypsin for overnight digestion. The next day, the samples were centrifuged at 16,000g for 20 min at room temperature to remove the NPs. The supernatant was acidified with TFA to a pH of 2-3 and cleaned using C18 StageTips. The samples were then heated at 95°C for 10 min, vacuum-dried, and submitted to the core facility for LC-MS analysis.

LC-MS/MS Analysis: Dried samples were reconstituted with 1 μg of peptides in 25 μl of LC loading buffer (3% ACN, 0.1% TFA) and analyzed using LC-MS/MS. A 60-min gradient was applied in LFQ mode, with 5 μl aliquots injected in triplicate. Control samples (55% human plasma) were prepared with 8 μg of peptides in 200 μl of loading buffer and analyzed similarly. An Ultimate 3000RSLCnano (Thermo Fisher) HPLC system was used with predefined columns, solvents, and gradient settings. Data Dependent Analysis (DDA) was performed with specific MS and MS2 scan settings, followed by data analysis using Proteome Discoverer 2.4 (Thermo Fisher), applying the protocols detailed in our earlier publication (center #9)^57^. The PtdChos concentration series experiment was performed using the same protocol, and the samples were analyzed over a 120 min gradient.

### Sample preparation for top-down proteomics

Protein elution from the surface of NPs and purification were conducted based on procedures illustrated in our recent publication^102^. The protein corona coated NPs (with/without PtdChos) were separately treated in a 0.4% (w/v) SDS solution at 60°C for 1.5 hours with continuous agitation to release the protein corona from the NP surface. Subsequently, the supernatant containing the protein corona in 0.4% SDS was separated from the NPs by centrifugation at 19,000xg for 20 min at 4°C. To ensure thorough separation, the supernatant underwent an additional centrifugation step under the same conditions. The final protein corona sample was then subjected to buffer exchange using an Amicon Ultra Centrifugal Filter with a 10 kDa Molecular Weight Cut-Off, effectively removing sodium dodecyl sulfate (SDS) from the protein samples.

The buffer exchange process began by wetting the filter with 20 µL of 100 mM ABC (pH 8.0), followed by centrifugation at 14,000xg for 10 min. Next, 200 µg of proteins were added to the filter, and centrifugation was conducted for 20 min at 14,000xg. This step was repeated with the addition of 200 µL of 8 M urea in 100 mM ammonium bicarbonate, followed by centrifugation for 20 min at 14,000xg, and repeated twice to ensure complete removal of SDS and other small molecules. To eliminate urea from the purified protein, the filter underwent three additional rounds of buffer exchange. Specifically, 100 mM ABC was added to the filter, adjusting the final volume to 200 µL. All procedures were carried out at 4°C to effectively eliminate urea from the protein corona.

Following buffer exchange, the total protein concentration was determined using a bicinchoninic acid (BCA) assay kit from Fisher Scientific (Hampton, NH), following the manufacturer’s instructions. The samples were then stored overnight at 4°C. The final protein solutions, consisting of 40 µL (without PtdChos initially) and 44 µL (with PtdChos initially) of 100 mM ABC with a protein concentration of 2.8 mg/mL, were prepared for LC-MS/MS analysis.

### Top-down proteomics LC-MS/MS

The RPLC separation was performed using an EASY-nLC™ 1200 system from Thermo Fisher Scientific. Each sample was diluted in 100 mM ABC. A 1 µL aliquot of the sample (0.3 mg/mL) was loaded onto a C4 pre-column (Acclaim PrepMapTM 100, 75 µm i.d. × 2 cm, nanoviper, 3 µm particles, 300 Å, Thermo Scientific). Subsequently, proteoforms were separated on a C4 analytical column (Acclaim PrepMapTM 100, 75 µm i.d. × 20 cm, nanoviper, 3 µm particles, 300 Å, Thermo Scientific) at a flow rate of 400 nL/min.A gradient composed of mobile phase A (2% ACN in water containing 0.1% FA) and mobile phase B (80% ACN with 0.1% FA) was used for separation. The gradient profile consisted of a 105-min program: 0-85 min, 8-70% B; 85-90 min, 70-100% B; 90-105 min, 100% B. The LC system required an additional 30 min for column equilibration between the analyses, resulting in approximately 135 min per LC-MS analysis.

The experiments utilized a Q-Exactive HF mass spectrometer, employing a data-dependent acquisition (DDA) method. MS settings included 120,000 mass resolution (at m/z 200), 3 microscans, a 3E6 AGC target value, maximum injection time of 100 ms, and a scan range of 600 to 2000 m/z. For MS/MS analysis, parameters included 120,000 mass resolution (at 200 m/z), 3 microscans, a 1E5 AGC target, 200 ms injection time, 4 m/z isolation window, and 20% normalized collision energy (NCE). During MS/MS, the top 5 most intense precursor ions from each MS spectrum were selected in the quadrupole and fragmented using higher-energy collision dissociation (HCD). Fragmentation occurred exclusively for ions with intensities exceeding 5E4 and charge states of 4 or higher. Dynamic exclusion was enabled with a 30-second duration, and the "Exclude isotopes" feature was activated.

### Top-down proteomics data analysis

Complex sample data were analyzed using Xcalibur software (Thermo Fisher Scientific) to obtain proteoform intensities and retention times. Chromatograms were exported from Xcalibur and formatted using Adobe Illustrator for final figure presentation.

Proteoform identification and quantification were conducted using the TopPIC Suite (Top-down mass spectrometry-based Proteoform Identification and Characterization, version 1.7.4) pipeline^103^. Initially, RAW files were converted to mzML format using the MSConvert tool. Spectral deconvolution, which converted precursor and fragment isotope clusters to monoisotopic masses, and proteoform feature detection were performed using TopFD (Top-down mass spectrometry Feature Detection, version 1.7.4)^104^. The resulting mass spectra were stored in msalign files, while proteoform feature information was stored in text files.

Database searches were carried out using TopPIC Suite against a custom-built protein database (∼2780 protein sequences), which included proteins identified in the BUP data. The search allowed for a maximum of one unexpected mass shift, with mass error tolerances of 10 ppm for precursors and fragments. Unknown mass shifts up to 500 Da were considered. False discovery rates (FDRs) for proteoform identifications were estimated using a target-decoy approach, filtering proteoform identifications at 1% and 5% FDR at the PrSM and proteoform levels, respectively.

Lists of identified proteoforms from all RPLC-MS/MS runs are provided in the **Supplementary Data 6**. Label-free quantification of identified proteoforms was performed using TopDiff (Top-down mass spectrometry-based identification of Differentially expressed proteoforms, version 1.7.4) with default settings^105^.

### LC-MS analysis by DIA

The samples were centrifuged at 14,000 g for 20 min to remove the unbound proteins. The collected NP pellets were washed three times with cold PBS under the same conditions. The samples were resuspended in 20 µl of PBS, and the proteins were reduced with 2 mM DTT (final concentration) for 45 min and then alkylated using 8 mM IAA (final concentration) for 45 min in the dark. Subsequently, 5 µl of LysC at 0.02 µg/µl was added for 4h, followed by the same concentration and volume of trypsin overnight. The samples were then centrifuged at 16,000g for 20 min at room temperature to remove the NPs then cleaned using C18 cartridges and vacuum dried.

Dried peptides were resuspended in 0.1% aqueous formic acid and subjected to LC–MS/MS analysis using an Exploris 480 Mass Spectrometer fitted with an Vanquish Neo (both Thermo Fisher Scientific) and a custom-made column heater set to 60°C. Peptides were resolved using a RP-HPLC column (75 μm × 30 cm) packed in-house with C18 resin (ReproSil-Pur C18–AQ, 1.9 μm resin; Dr. Maisch GmbH) at a flow rate of 0.2 μL/min. The following gradient was used for peptide separation: from 4% B to 10% B over 7.5 min to 35% B over 67.5 min to 50% B over 15 min to 95% B over 1 min followed by 10 min at 95% B to 5% B over 1 min followed by 4 min at 5% B. Buffer A was 0.1% formic acid in water and buffer B was 80% acetonitrile, 0.1% formic acid in water.

The mass spectrometer was operated in DIA mode with a cycle time of 3 seconds. MS1 scans were acquired in the Orbitrap in centroid mode at a resolution of 120,000 FWHM (at 200 m/z), a scan range from 390 to 910 m/z, normalized AGC target set to 300 % and maximum ion injection time mode set to Auto. MS2 scans were acquired in the Orbitrap in centroid mode at a resolution of 15,000 FWHM (at 200 m/z), precursor mass range of 400 to 900, quadrupole isolation window of 7 m/z with 1 m/z window overlap, a defined first mass of 120 m/z, normalized AGC target set to 3000% and a maximum injection time of 22 ms. Peptides were fragmented by HCD with collision energy set to 28% and one microscan was acquired for each spectrum.

The acquired RAW files were searched individually using the Spectronaut (Biognosys v18.6) directDIA workflow against a *Homo sapiens* database (consisting of 20,360 protein sequences downloaded from Uniprot on 2022/02/22) and 392 commonly observed contaminants. Default settings were used.

For analysis of the impact of PtdChos treated plasma and different NPs, we chose a quicker LC-MS setup (30 SPD) consisting of an Exploris 480 fitted with an Evosep One using the following settings.

Dried peptides were resuspended in 0.1% aqueous formic acid, loaded onto Evotip Pure tips (Evosep Biosystems) and subjected to LC–MS/MS analysis using an Exploris 480 Mass Spectrometer (Thermo Fisher Scientific) fitted with an Evosep One (EV 1000, Evosep Biosystems). Peptides were resolved using a Performance Column – 30 SPD (150 μm × 15 cm, 1.5 um, EV1137, Evosep Biosystems) kept at 40 °C fitted with a stainless-steel emitter (30 um, EV1086, Evosep Biosystems) using the 30 SPD method. Buffer A was 0.1% formic acid in water and buffer B was acetonitrile, with 0.1% formic acid.

The mass spectrometer was operated in DIA mode. MS1 scans were acquired in centroid mode at a resolution of 120,000 FWHM (at 200 m/z), a scan range from 350 to 1500 m/z, AGC target set to standard and maximum ion injection time mode set to Auto. MS2 scans were acquired in centroid mode at a resolution of 15,000 FWHM (at 200 m/z), precursor mass range of 400 to 900 m/z, quadrupole isolation window of 12 m/z without window overlap, a defined first mass of 120 m/z, normalized AGC target set to 3000 % and maximum injection time mode set to Auto. Peptides were fragmented by HCD with collision energy set to 28% and one microscan was acquired for each spectrum.

The acquired RAW files were searched using the Spectronaut (Biognosys v19.0) directDIA workflow against a *Homo sapiens* database (consisting of 20,360 protein sequences downloaded from Uniprot on 2022/02/22) and 392 commonly observed contaminants. Default settings were applied except Method Evaluation was set to TRUE.

### *In silico* experiments

The crystal structure of human serum albumin (PDB Code: 1AO6) was obtained from the protein data bank and used for all simulation setups (**Supplementary Fig. 11**). The structure of the PtdChos ligand was obtained from the CHARMM 36 force field files (Name: PLPC).^106^

### Molecular Docking

Two blind docking methods and one site specific docking were performed with Autodock Vina^107, 108^ software. The first blind docking used the whole albumin for the binding search and the second method consisted of multiple search boxes covering the entire albumin surface. The site-specific docking was performed based on crystallographic analysis of the binding sites on albumin for palmitic acid.^109^ The top 10 unique non-overlapping binding poses were kept for the subsequent molecular dynamics simulations.

### MD simulations

All-atom MD simulations were performed with GROMACS^110^ free software and the CHARMM36^111^ force field. Four types of protein-ligand systems were investigated (**Supplementary Fig. 11 b-c**). Three 1 ligand systems, one 3 ligands, 5 ligands, and 10 ligands systems each were used for the simulations. The protein-ligand systems along with TIP3P water model and a neutralizing salt-concentration of 0.15 M NaCl were energy minimized using 5000 steps with an energy tolerance of 1000 KJ/mol/nm. The systems were subsequently equilibrated in a 1 ns NVT and 4 ns NPT steps with a 1 fs timestep. The constant temperature for all runs was 310 K and the Berendsen pressure coupling was used. Production steps were then run for 100 ns with a 2 fs with the with the Parrinello-Rahman barostat.

### Post-processing analysis

#### Interaction energy

The short range nonbonded Coulombic and Lennard-Jones interaction energies between albumin and the ligands were calculated using GROMACS^110^.

#### Free energy calculation

The free energy calculation for the entire 100 ns simulations were calculated using the gmx_MMPBSA^112, 113^ package with the generalized Born method. The entropic term was not considered. The residues and ligand atoms within 6 Angstroms were selected for the calculation. The gas phase and solvation terms were averaged for the 1 ligand systems and plotted alongside the 3, 5, and 10 ligand systems.

#### RMSF

Root mean square fluctuation per residue was calculated using GROMACS^110^ free software after fitting to the first frame of the simulation. For the 1 ligand systems, all 3 simulations results were averaged.

#### RMSD

Root mean square deviation of the ligand(s) with respect to the energy minimized structure was calculated using GROMACS^110^. The results of the three 1 ligand systems were averaged.

#### Bond types

The types of bonds formed between albumin and PtdChos were determined using the MD-Ligand-Receptor tool^114^.

#### Visualization

All visualizations were made using the Visual Molecular Dynamics (VMD) software^115^.

## Data analysis

First, data were normalized by total protein intensity in each technical replicate. Then all the abundances were transformed into log10 and NA values were imputed by a constant value of –10 (in the heatmap figure). Except for PtdChos sample at 100 µg/ml, all samples were analyzed with three technical replicates. In case of DIA analysis of different nanoparticles, there were 4 individual samples per group with no technical replicates. Statistcal T-test with unequal variance were used to compare the differences between groups. Data analysis was performed using R (R version 4.1.0) with the help of ggplot2, dplyr, tidyr, ComplexHeatmap and PerfromanceAnalytics packages.

## Statistics and reproducibility

All measurements were performed as a triplicate analysis of a given aliquot. The initial DIA analysis was performed in one replicate. The experiments on different NPs with PtdChos and DIA were performed on plasma samples from four individual donors.

## Data availability

The authors declare that all data supporting the findings of this study are available within the paper and its supplementary information and data files. The mass spectrometry data for all the bottom-up experiments are deposited in the database MassIVE with the identifiers MSV000094257. The MS RAW files for the top-down proteomics analysis were submitted to the ProteomeXchange Consortium through PRIDE^116^ and assigned the dataset identifier PXD053359. All data are also available from the corresponding authors (A.A.S. and M.M.)

## Materials & Correspondence

Corresponding authors: A.A.S. and (M.M.)

## Acknowledgement

MM gratefully acknowledge financial support from the U.S. National Institute of Diabetes and Digestive and Kidney Diseases (grant DK131417). A.A.S. was supported by an Ambizione Fellowship from the Swiss National Science Foundation (SNSF grant number: PZ00P3_216203) and a grant from Karolinska Institutet (2-188/2022). L.S. thanks the support from the National Cancer Institute (NCI) through the grant R01CA247863. We acknowledge support of a Burroughs Wellcome Fund Career Award at the Scientific Interface (CASI) (MPL), a Dreyfus foundation award (MPL), the Philomathia foundation (MPL), an NSF CAREER award 2046159 (MPL), an NSF CBET award 1733575 (to MPL), a CZI imaging award (MPL), a Sloan Foundation Award (MPL), a McKnight Foundation award (MPL), a Simons Foundation Award (MPL), a Moore Foundation Award (MPL), a Brain Foundation Award (MPL), and a polymaths award from Schmidt Sciences, LLC (MPL). MPL is a Chan Zuckerberg Biohub San Francisco investigator.

## References

1. Schwenk JM, et al. The human plasma proteome draft of 2017: building on the human plasma PeptideAtlas from mass spectrometry and complementary assays. Journal of proteome research 16, 4299–4310 (2017).

2. Zubarev RA. The challenge of the proteome dynamic range and its implications for in-depth proteomics. Proteomics 13, 723–726 (2013).

3. Zhang Q, Faca V, Hanash S. Mining the plasma proteome for disease applications across seven logs of protein abundance. Journal of proteome research 10, 46–50 (2011).

4. Pernemalm M, et al. In-depth human plasma proteome analysis captures tissue proteins and transfer of protein variants across the placenta. Elife 8, e41608 (2019).

5. Blume JE, et al. Rapid, deep and precise profiling of the plasma proteome with multi-nanoparticle protein corona. Nature Communications 11, 3662 (2020).

6. Zhu G, et al. Single Chain Variable Fragment Displaying M13 Phage Library Functionalized Magnetic Microsphere-Based Protein Equalizer for Human Serum Protein Analysis. Analytical Chemistry 84, 7633–7637 (2012).

7. Fonslow BR, et al. Digestion and depletion of abundant proteins improves proteomic coverage. Nature Methods 10, 54–56 (2013).

8. Geyer PE, et al. Plasma Proteome Profiling to detect and avoid sample-related biases in biomarker studies. EMBO molecular medicine 11, e10427 (2019).

9. Ignjatovic V, et al. Mass spectrometry-based plasma proteomics: considerations from sample collection to achieving translational data. Journal of proteome research 18, 4085–4097 (2019).

10. Pattipeiluhu R, Crielaard S, Klein-Schiphorst I, Florea BI, Kros A, Campbell F. Unbiased Identification of the Liposome Protein Corona using Photoaffinity-based Chemoproteomics. ACS Central Science 6, 535–545 (2020).

11. Saei AA, et al. ProTargetMiner as a proteome signature library of anticancer molecules for functional discovery. Nature communications 10, 5715 (2019).

12. Woo J, Zhang Q. A Streamlined High-Throughput Plasma Proteomics Platform for Clinical Proteomics with Improved Proteome Coverage, Reproducibility, and Robustness. Journal of the American Society for Mass Spectrometry 34, 754–762 (2023).

13. Viode A, et al. A simple, time- and cost-effective, high-throughput depletion strategy for deep plasma proteomics. Science Advances 9, eadf9717 (2023).

14. Palstrøm NB, Rasmussen LM, Beck HC. Affinity capture enrichment versus affinity depletion: a comparison of strategies for increasing coverage of low-abundant human plasma proteins. International journal of molecular sciences 21, 5903 (2020).

15. Pringels L, Broeckx V, Boonen K, Landuyt B, Schoofs L. Abundant plasma protein depletion using ammonium sulfate precipitation and Protein A affinity chromatography. Journal of Chromatography B 1089, 43–59 (2018).

16. Tu C, et al. Depletion of abundant plasma proteins and limitations of plasma proteomics. J Proteome Res 9, 4982–4991 (2010).

17. Hadjidemetriou M, Al-Ahmady Z, Buggio M, Swift J, Kostarelos K. A novel scavenging tool for cancer biomarker discovery based on the blood-circulating nanoparticle protein corona. Biomaterials 188, 118–129 (2019).

18. Papafilippou L, Claxton A, Dark P, Kostarelos K, Hadjidemetriou M. Protein corona fingerprinting to differentiate sepsis from non-infectious systemic inflammation. Nanoscale 12, 10240–10253 (2020).

19. Papafilippou L, Claxton A, Dark P, Kostarelos K, Hadjidemetriou M. Nanotools for sepsis diagnosis and treatment. Advanced Healthcare Materials 10, 2001378 (2021).

20. Monopoli MP, et al. Physical-Chemical aspects of protein corona: Relevance to in vitro and in vivo biological impacts of nanoparticles. Journal of the American Chemical Society 133, 2525–2534 (2011).

21. Meng Y, et al. A highly efficient protein corona-based proteomic analysis strategy for the discovery of pharmacodynamic biomarkers. J Pharm Anal 12, 879–888 (2022).

22. Caracciolo G, et al. Disease-specific protein corona sensor arrays may have disease detection capacity. Nanoscale Horizons 4, 1063–1076 (2019).

23. Jiang Y, Meyer JG. Rapid Plasma Proteome Profiling via Nanoparticle Protein Corona and Direct Infusion Mass Spectrometry. Journal of Proteome Research 23, 3649–3658 (2024).

24. Mekseriwattana W, Thiangtrongjit T, Reamtong O, Wongtrakoongate P, Katewongsa KP. Proteomic Analysis Reveals Distinct Protein Corona Compositions of Citrate- and Riboflavin-Coated SPIONs. ACS Omega 7, 37589–37599 (2022).

25. Qiu L, et al. How eluents define proteomic fingerprinting of protein corona on nanoparticles. Journal of Colloid and Interface Science 648, 497–510 (2023).

26. Caracciolo G, et al. Disease-specific protein corona sensor arrays may have disease detection capacity. Nanoscale Horizons 4, 1063–1076 (2019).

27. Mahmoudi M, Landry MP, Moore A, Coreas R. The protein corona from nanomedicine to environmental science. Nature Reviews Materials 8, 422–438 (2023).

28. Hajipour MJ, et al. An Overview of Nanoparticle Protein Corona Literature. Small n/a, 2301838.

29. Jiang Y, Meyer JG. 1.4 min Plasma Proteome Profiling via Nanoparticle Protein Corona and Direct Infusion Mass Spectrometry. bioRxiv, 2024.2002. 2006.579213 (2024).

30. Sharifi S, Reuel N, Kallmyer N, Sun E, Landry MP, Mahmoudi M. The Issue of Reliability and Repeatability of Analytical Measurement in Industrial and Academic Nanomedicine. ACS nano 17, 4–11 (2022).

31. Tang H, et al. Cholesterol modulates the physiological response to nanoparticles by changing the composition of protein corona. Nature Nanotechnology, (2023).

32. Mahmoudi N, Mahmoudi M. Effects of cholesterol on biomolecular corona. Nature Nanotechnology, (2023).

33. Fonda ML. Vitamin B6 metabolism and binding to proteins in the blood of alcoholic and nonalcoholic men. Alcoholism: Clinical and Experimental Research 17, 1171–1178 (1993).

34. Panja S, Khatua DK, Halder M. Simultaneous Binding of Folic Acid and Methotrexate to Human Serum Albumin: Insights into the Structural Changes of Protein and the Location and Competitive Displacement of Drugs. ACS Omega 3, 246–253 (2018).

35. Ghosh R, Thomas DS, Arcot J. Molecular Recognition Patterns between Vitamin B12 and Proteins Explored through STD-NMR and In Silico Studies. Foods 12, 575 (2023).

36. Jackowski S, Alix J-H. Cloning, sequence, and expression of the pantothenate permease (panF) gene of Escherichia coli. Journal of bacteriology 172, 3842–3848 (1990).

37. Musa TL, Ioerger TR, Sacchettini JC. The Tuberculosis Structural Genomics Consortium: A Structural Genomics Approachto Drug Discovery. In: Advances in Protein Chemistry and Structural Biology (ed Joachimiak A). Academic Press (2009).

38. Adhel E, Ha Duong N-T, Vu TH, Taverna D, Ammar S, Serradji N. Interaction between carbon dots from folic acid and their cellular receptor: a qualitative physicochemical approach. Physical Chemistry Chemical Physics 25, 14324–14333 (2023).

39. Lonsdale D. Thiamin and protein folding. Medical Hypotheses 129, 109252 (2019).

40. Mkrtchyan G, et al. Molecular mechanisms of the non-coenzyme action of thiamin in brain: biochemical, structural and pathway analysis. Scientific Reports 5, 12583 (2015).

41. Lee DW, Park YW, Lee MY, Jeong KH, Lee JY. Structural analysis and insight into effector binding of the niacin-responsive repressor NiaR from Bacillus halodurans. Scientific Reports 10, 21039 (2020).

42. Mahmoudi M, Lohse SE, Murphy CJ, Fathizadeh A, Montazeri A, Suslick KS. Variation of protein corona composition of gold nanoparticles following plasmonic heating. Nano letters 14, 6–12 (2013).

43. Saha K, et al. Regulation of Macrophage Recognition through the Interplay of Nanoparticle Surface Functionality and Protein Corona. ACS nano 10, 4421–4430 (2016).

44. Ashkarran AA, et al. Protein Corona Composition of Gold Nanocatalysts. ACS Pharmacology & Translational Science 7, 1169–1177 (2024).

45. Askim JR, Mahmoudi M, Suslick KS. Optical sensor arrays for chemical sensing: the optoelectronic nose. Chemical Society Reviews 42, 8649–8682 (2013).

46. Mahmoudi M, et al. Irreversible changes in protein conformation due to interaction with superparamagnetic iron oxide nanoparticles. Nanoscale 3, 1127–1138 (2011).

47. Sakulkhu U, Mahmoudi M, Maurizi L, Salaklang J, Hofmann H. Protein corona composition of superparamagnetic iron oxide nanoparticles with various physico-chemical properties and coatings. Scientific reports 4, 5020 (2014).

48. Mahmoudi M, Akhavan O, Ghavami M, Rezaee F, Ghiasi SMA. Graphene oxide strongly inhibits amyloid beta fibrillation. Nanoscale 4, 7322–7325 (2012).

49. Mao H, et al. Hard corona composition and cellular toxicities of the graphene sheets. Colloids and Surfaces B: Biointerfaces 109, 212–218 (2013).

50. Hajipour MJ, et al. Personalized disease-specific protein corona influences the therapeutic impact of graphene oxide. Nanoscale 7, 8978–8994 (2015).

51. Rahman M, Laurent S, Tawil N, Yahia LH, Mahmoudi M. Protein-nanoparticle interactions: the bio-nano interface.). Springer Science & Business Media (2013).

52. Rahimi M, et al. Zeolite Nanoparticles for Selective Sorption of Plasma Proteins. Scientific reports 5, 17259–17259 (2014).

53. Laurent S, et al. Corona protein composition and cytotoxicity evaluation of ultra-small zeolites synthesized from template free precursor suspensions. Toxicology Research 2, 270–279 (2013).

54. Hajipour MJ, Laurent S, Aghaie A, Rezaee F, Mahmoudi M. Personalized protein coronas: a “key” factor at the nanobiointerface. Biomaterials science 2, 1210–1221 (2014).

55. Ashkarran AA, et al. Sex-Specific Silica Nanoparticle Protein Corona Compositions Exposed to Male and Female BALB/c Mice Plasmas. ACS Bio & Med Chem Au 3, 62–73 (2023).

56. Sheibani S, et al. Nanoscale characterization of the biomolecular corona by cryo-electron microscopy, cryo-electron tomography, and image simulation. Nature communications 12, 573 (2021).

57. Ashkarran AA, Gharibi H, Voke E, Landry MP, Saei AA, Mahmoudi M. Measurements of heterogeneity in proteomics analysis of the nanoparticle protein corona across core facilities. Nature Communications 13, 6610 (2022).

58. Gharibi H, et al. A uniform data processing pipeline enables harmonized nanoparticle protein corona analysis across proteomics core facilities. Nature Communications 15, 342 (2024).

59. Ashkarran AA, Ghavami M, Aghaverdi H, Stroeve P, Mahmoudi M. Bacterial effects and protein corona evaluations: crucial ignored factors in the prediction of bio-efficacy of various forms of silver nanoparticles. Chemical research in toxicology 25, 1231–1242 (2012).

60. Bigdeli A, et al. Exploring Cellular Interactions of Liposomes Using Protein Corona Fingerprints and Physicochemical Properties. ACS nano 10, 3723–3737 (2016).

61. Palchetti S, et al. Nanoparticles-cell association predicted by protein corona fingerprints. Nanoscale 8, 12755–12763 (2016).

62. Lunov O, et al. Differential Uptake of Functionalized Polystyrene Nanoparticles by Human Macrophages and a Monocytic Cell Line. ACS Nano 5, 1657–1669 (2011).

63. Tenzer S, et al. Nanoparticle Size Is a Critical Physicochemical Determinant of the Human Blood Plasma Corona: A Comprehensive Quantitative Proteomic Analysis. ACS Nano 5, 7155–7167 (2011).

64. Tonigold M, et al. Pre-adsorption of antibodies enables targeting of nanocarriers despite a biomolecular corona. Nature Nanotechnology 13, 862–869 (2018).

65. Monopoli MP, Åberg C, Salvati A, Dawson KA. Biomolecular coronas provide the biological identity of nanosized materials. Nature Nanotechnology 7, 779–786 (2012).

66. Tenzer S, et al. Rapid formation of plasma protein corona critically affects nanoparticle pathophysiology. Nature Nanotechnology 8, 772–781 (2013).

67. Sheibani S, et al. Nanoscale characterization of the biomolecular corona by cryo-electron microscopy, cryo-electron tomography, and image simulation. Nature Communications 12, 573 (2021).

68. Mahmoudi M. The need for improved methodology in protein corona analysis. Nature Communications 13, 49 (2022).

69. Tavakol M, et al. Disease-related metabolites affect protein–nanoparticle interactions. Nanoscale 10, 7108–7115 (2018).

70. Hamilton JA. Interactions of triglycerides with phospholipids: incorporation into the bilayer structure and formation of emulsions. Biochemistry 28, 2514–2520 (1989).

71. Saito H, Tanaka M, Okamura E, Kimura T, Nakahara M, Handa T. Interactions of Phosphatidylcholine Surface Monolayers with Triglyceride Cores and Enhanced ApoA-1 Binding in Lipid Emulsions. Langmuir 17, 2528–2532 (2001).

72. Cassidy L, Kaulich PT, Maaß S, Bartel J, Becher D, Tholey A. Bottom-up and top-down proteomic approaches for the identification, characterization, and quantification of the low molecular weight proteome with focus on short open reading frame-encoded peptides. PROTEOMICS 21, 2100008 (2021).

73. Picotti P, Aebersold R. Selected reaction monitoring-based proteomics: workflows, potential, pitfalls and future directions. Nat Methods 9, 555–566 (2012).

74. Roberts DS, et al. Top-down proteomics. Nat Rev Methods Primers 4, (2024).

75. Yates JR, Ruse CI, Nakorchevsky A. Proteomics by Mass Spectrometry: Approaches, Advances, and Applications. Annual Review of Biomedical Engineering 11, 49–79 (2009).

76. Ashkarran AA, Gharibi H, Modaresi SM, Saei AA, Mahmoudi M. Standardizing Protein Corona Characterization in Nanomedicine: A Multicenter Study to Enhance Reproducibility and Data Homogeneity. Nano Letters 24, 9874–9881 (2024).

77. Tenzer S, et al. Rapid formation of plasma protein corona critically affects nanoparticle pathophysiology. Nature nanotechnology 8, 772 (2013).

78. Mahley RW. Apolipoprotein E: from cardiovascular disease to neurodegenerative disorders. Journal of molecular medicine 94, 739–746 (2016).

79. Mahmoudi N, Mahmoudi M. Effects of cholesterol on biomolecular corona. Nature Nanotechnology 18, 974–976 (2023).

80. Lundqvist M, Stigler J, Elia G, Lynch I, Cedervall T, Dawson KA. Nanoparticle size and surface properties determine the protein corona with possible implications for biological impacts. Proceedings of the National Academy of Sciences 105, 14265–14270 (2008).

81. Monopoli MP, et al. Physical−Chemical Aspects of Protein Corona: Relevance to in Vitro and in Vivo Biological Impacts of Nanoparticles. Journal of the American Chemical Society 133, 2525–2534 (2011).

82. Walczyk D, Bombelli FB, Monopoli MP, Lynch I, Dawson KA. What the Cell “Sees” in Bionanoscience. Journal of the American Chemical Society 132, 5761–5768 (2010).

83. Lesniak A, Fenaroli F, Monopoli MP, Åberg C, Dawson KA, Salvati A. Effects of the Presence or Absence of a Protein Corona on Silica Nanoparticle Uptake and Impact on Cells. ACS Nano 6, 5845–5857 (2012).

84. Ghavami M, et al. Plasma concentration gradient influences the protein corona decoration on nanoparticles. RSC Advances 3, 1119–1126 (2013).

85. Mahmoudi M, et al. The protein corona mediates the impact of nanomaterials and slows amyloid beta fibrillation. ChemBioChem 14, 568–572 (2013).

86. Aebersold R, et al. How many human proteoforms are there? Nature chemical biology 14, 206–214 (2018).

87. Smith LM, Kelleher NL. Proteoforms as the next proteomics currency. Science 359, 1106–1107 (2018).

88. McCool EN, et al. Deep top-down proteomics revealed significant proteoform-level differences between metastatic and nonmetastatic colorectal cancer cells. Science Advances 8, eabq6348 (2022).

89. Tucholski T, et al. Distinct hypertrophic cardiomyopathy genotypes result in convergent sarcomeric proteoform profiles revealed by top-down proteomics. Proceedings of the National Academy of Sciences 117, 24691–24700 (2020).

90. Sheibani S, et al. Nanoscale Characterization of the Biomolecular Corona by Cryo-Electron Microscopy, Cryo-Electron Tomography, and Image Simulation. Nature Communications 12, 573 (2021).

91. Biterova EI, et al. The crystal structure of human microsomal triglyceride transfer protein. Proceedings of the National Academy of Sciences 116, 17251–17260 (2019).

92. Volanakis JE, Wirtz KW. Interaction of C-reactive protein with artificial phosphatidylcholine bilayers. Nature 281, 155–157 (1979).

93. Sengupta T, Manoj N. Phosphatidylserine and phosphatidylethanolamine bind to protein Z cooperatively and with equal affinity. PLoS One 11, e0161896 (2016).

94. Boggs JM, Rangaraj G, Dicko A. Effect of phosphorylation of phosphatidylinositol on myelin basic protein-mediated binding of actin filaments to lipid bilayers in vitro. Biochimica et Biophysica Acta (BBA) - Biomembranes 1818, 2217–2227 (2012).

95. Roderick SL, et al. Structure of human phosphatidylcholine transfer protein in complex with its ligand. Nature Structural Biology 9, 507–511 (2002).

96. Jonas A. Interaction of phosphatidylcholine with bovine serum albumin. Specificity and properties of the complexes. Biochimica et Biophysica Acta (BBA)-Protein Structure 427, 325–336 (1976).

97. Zborowski J, Roerdink F, Scherphof G. Leakage of sucrose from phosphatidylcholine liposomes induced by interaction with serum albumin. Biochimica et Biophysica Acta (BBA)-General Subjects 497, 183–191 (1977).

98. Morrisett J, Jackson R, Gotto Jr A. Lipid-protein interactions in the plasma lipoproteins. Biochimica et Biophysica Acta (BBA)-Reviews on Biomembranes 472, 93–133 (1977).

99. Adams LM, et al. Mapping the KRAS proteoform landscape in colorectal cancer identifies truncated KRAS4B that decreases MAPK signaling. J Biol Chem 299, 102768 (2023).

100. Toby TK, Fornelli L, Kelleher NL. Progress in Top-Down Proteomics and the Analysis of Proteoforms. Annu Rev Anal Chem (Palo Alto Calif) 9, 499–519 (2016).

101. Shi T, et al. IgY14 and SuperMix immunoaffinity separations coupled with liquid chromatography-mass spectrometry for human plasma proteomics biomarker discovery. Methods 56, 246–253 (2012).

102. Sadeghi SA, Ashkarran AA, Mahmoudi M, Sun L. Mass spectrometry-based top-down proteomics in nanomedicine: proteoform-specific measurement of protein corona. bioRxiv, 2024.2003.2022.586273 (2024).

103. Duchêne S, Geoghegan JL, Holmes EC, Ho SYW. Estimating evolutionary rates using time-structured data: a general comparison of phylogenetic methods. Bioinformatics 32, 3375–3379 (2016).

104. Basharat AR, Zang Y, Sun L, Liu X. TopFD: A Proteoform Feature Detection Tool for Top–Down Proteomics. Analytical Chemistry 95, 8189–8196 (2023).

105. Lubeckyj RA, Basharat AR, Shen X, Liu X, Sun L. Large-Scale Qualitative and Quantitative Top-Down Proteomics Using Capillary Zone Electrophoresis-Electrospray Ionization-Tandem Mass Spectrometry with Nanograms of Proteome Samples. Journal of the American Society for Mass Spectrometry 30, 1435–1445 (2019).

106. Klauda JB, et al. Update of the CHARMM All-Atom Additive Force Field for Lipids: Validation on Six Lipid Types. The Journal of Physical Chemistry B 114, 7830–7843 (2010).

107. Trott O, Olson AJ. AutoDock Vina: improving the speed and accuracy of docking with a new scoring function, efficient optimization, and multithreading. J Comput Chem 31, 455–461 (2010).

108. Eberhardt J, Santos-Martins D, Tillack AF, Forli S. AutoDock Vina 1.2.0: New Docking Methods, Expanded Force Field, and Python Bindings. Journal of Chemical Information and Modeling 61, 3891-3898 (2021).

109. Bhattacharya AA, Grüne T, Curry S. Crystallographic analysis reveals common modes of binding of medium and long-chain fatty acids to human serum albumin. J Mol Biol 303, 721–732 (2000).

110. Pronk S, et al. GROMACS 4.5: a high-throughput and highly parallel open source molecular simulation toolkit. Bioinformatics 29, 845–854 (2013).

111. Best RB, et al. Optimization of the Additive CHARMM All-Atom Protein Force Field Targeting Improved Sampling of the Backbone 1, ψ and Side-Chain χ1 and χ2 Dihedral Angles. Journal of Chemical Theory and Computation 8, 3257-3273 (2012).

112. Valdés-Tresanco MS, Valdés-Tresanco ME, Valiente PA, Moreno E. gmx_MMPBSA: A New Tool to Perform End-State Free Energy Calculations with GROMACS. Journal of Chemical Theory and Computation 17, 6281–6291 (2021).

113. Miller BR, III, McGee TD, Jr., Swails JM, Homeyer N, Gohlke H, Roitberg AE. MMPBSA.py: An Efficient Program for End-State Free Energy Calculations. Journal of Chemical Theory and Computation 8, 3314–3321 (2012).

114. Pieroni M, et al. MD-Ligand-Receptor: A High-Performance Computing Tool for Characterizing Ligand-Receptor Binding Interactions in Molecular Dynamics Trajectories. Int J Mol Sci 24, (2023).

115. Humphrey W, Dalke A, Schulten K. VMD: visual molecular dynamics. J Mol Graph 14, 33–38, 27-38 (1996).

116. Perez-Riverol Y, et al. The PRIDE database and related tools and resources in 2019: improving support for quantification data. Nucleic Acids Res 47, D442–d450 (2019).

