## Supplementary material for "Deep Plasma Proteome Profiling by Modulating Single Nanoparticle Protein Corona with Small Molecules": Supp Information

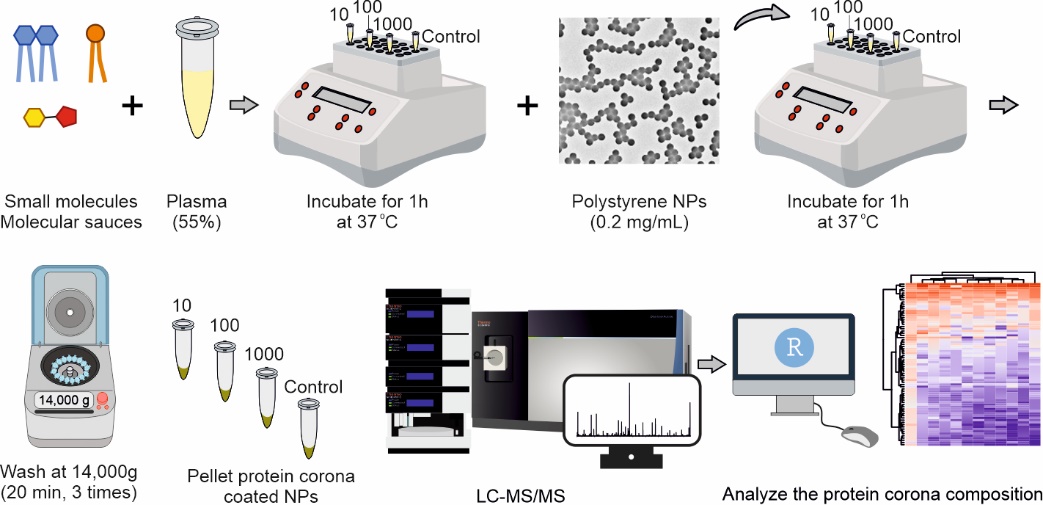

**Supplementary Scheme 1. The overall workflow of the study.** After exposing small molecules to human plasma, NPs were incubated with the treated or untreated plasma, purified, isolated, and used for analysis of the protein corona profile on the surface of the NPs using bottom-up and/or top-down LC-MS/MS.

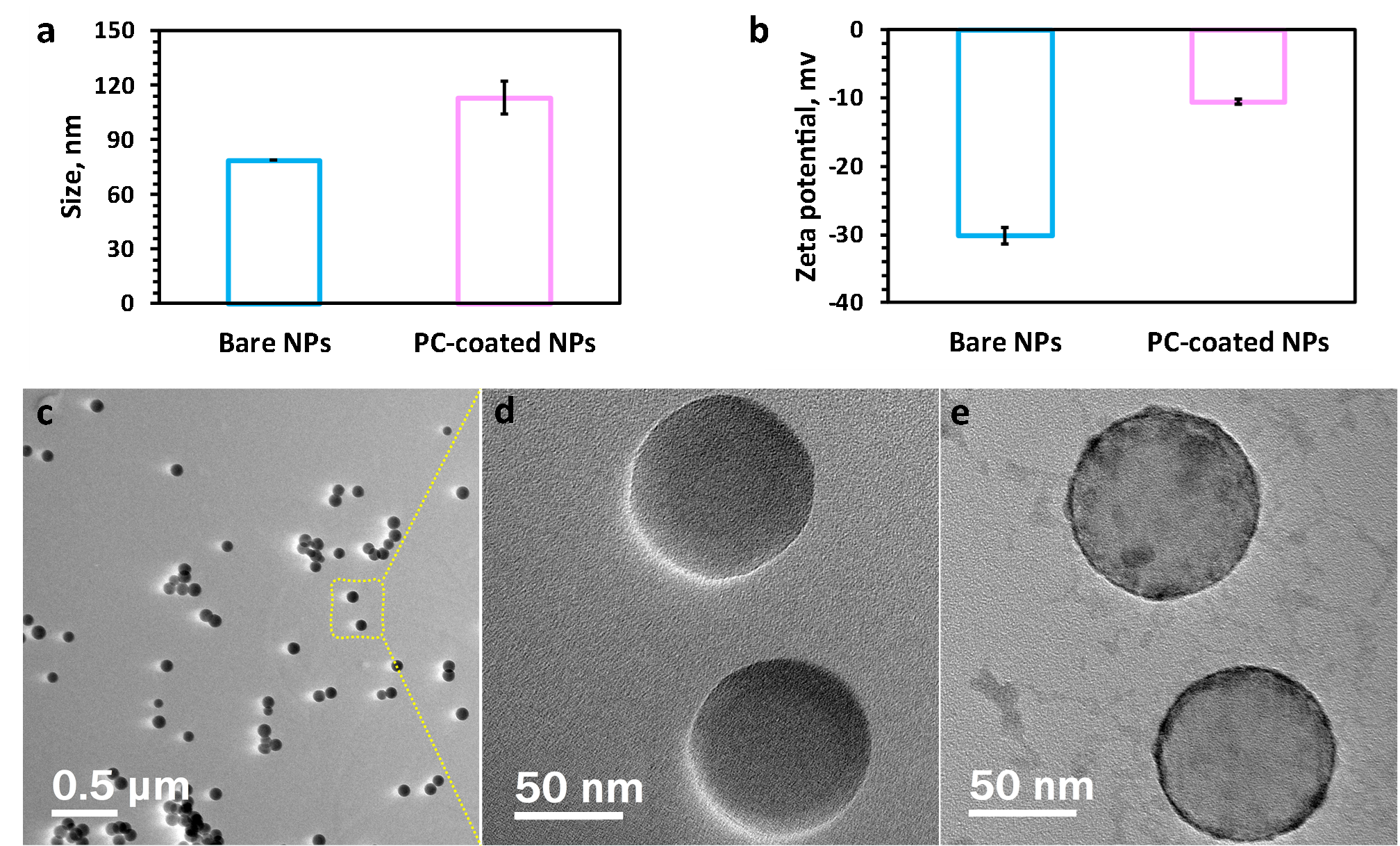

**Supplementary Fig. 1. Characterizations of the bare NPs and untreated protein corona coated NPs.** (**a**) and (**b**), DLS and zeta potential analysis of bare NPs and untreated protein corona–coated NPs respectively, (**c**), and (**d**) TEM images of bare polystyrene NPs, and (**e**) TEM image of protein corona–coated NPs, as representative. The polydispersity index (PDI) of bare and protein corona-coated NPs were found to be 0.023 and 0.214, respectively.

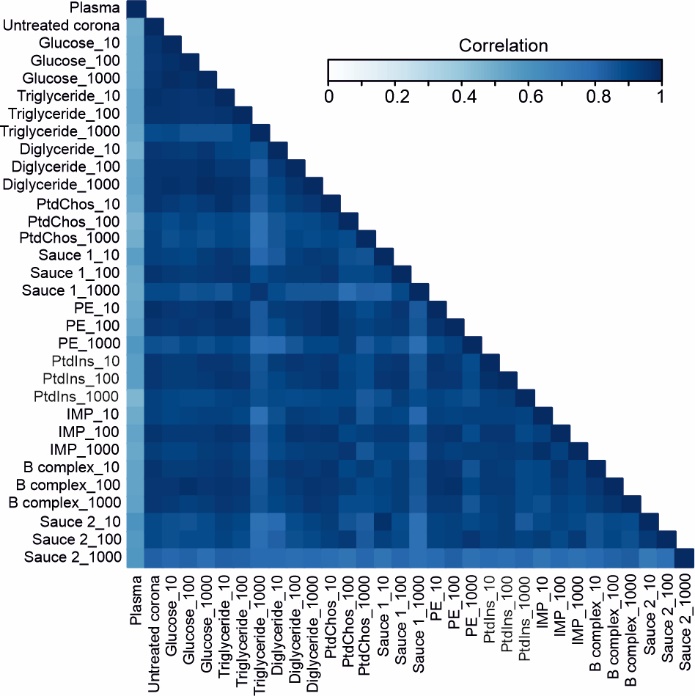

**Supplementary Fig. 2. Correlation of plasma proteome profiles**. Pearson correlation of the 117 shared proteins across all the samples (10-1000 µg/ml).

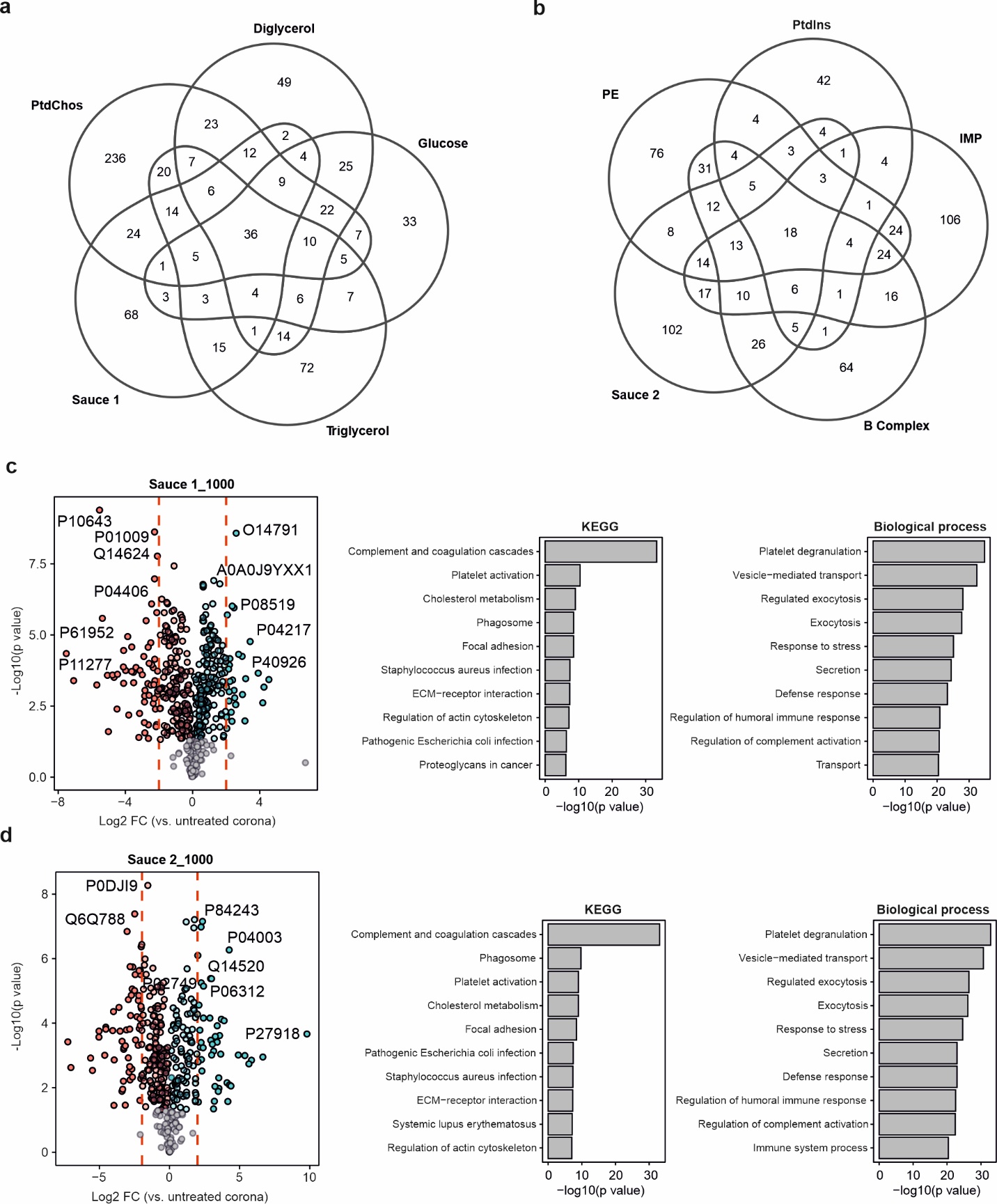

**Supplementary Fig. 3. The molecular sauces enrich or deplete specific proteins**. **a-b,** The number of unique proteins that were quantified in a given group which were not quantified in the plasma or with bare NPs. **c-d,** The enriched and depleted proteins for molecular sauce 1 and 2 in comparison to the untreated protein corona are shown (left panels). Respective pathway analysis was performed for all the depleted and enriched proteins (right panels). Significance was calculated using Student’s t-test.

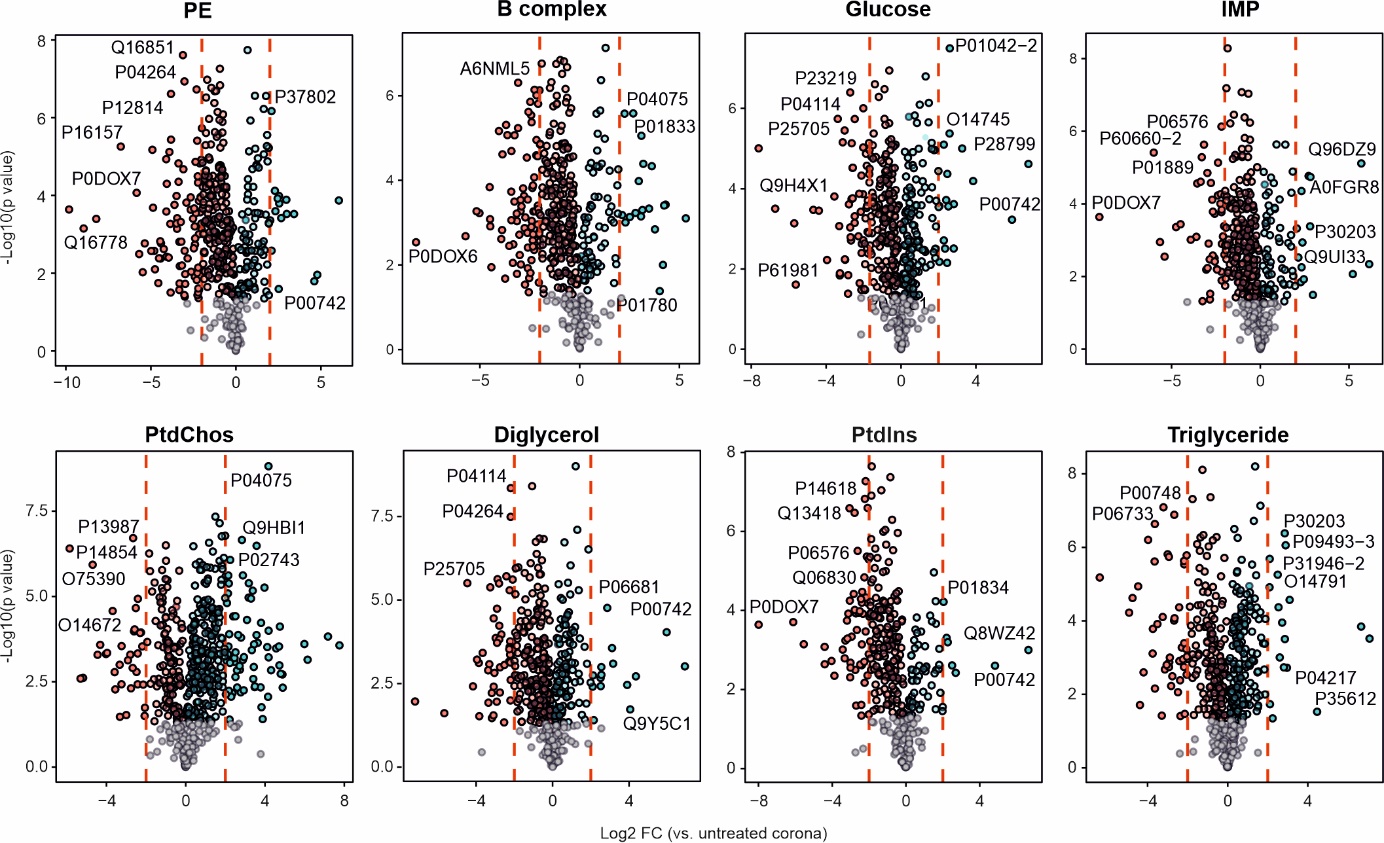

**Supplementary Fig. 4. The enrichment and depletion of specific proteins (only those shared) by spiking small molecules in NP protein corona vs. the abundance of proteins in the untreated NP protein corona.** Only the results for the highest concentration of each small molecule (1000 µg/ml) are shown.

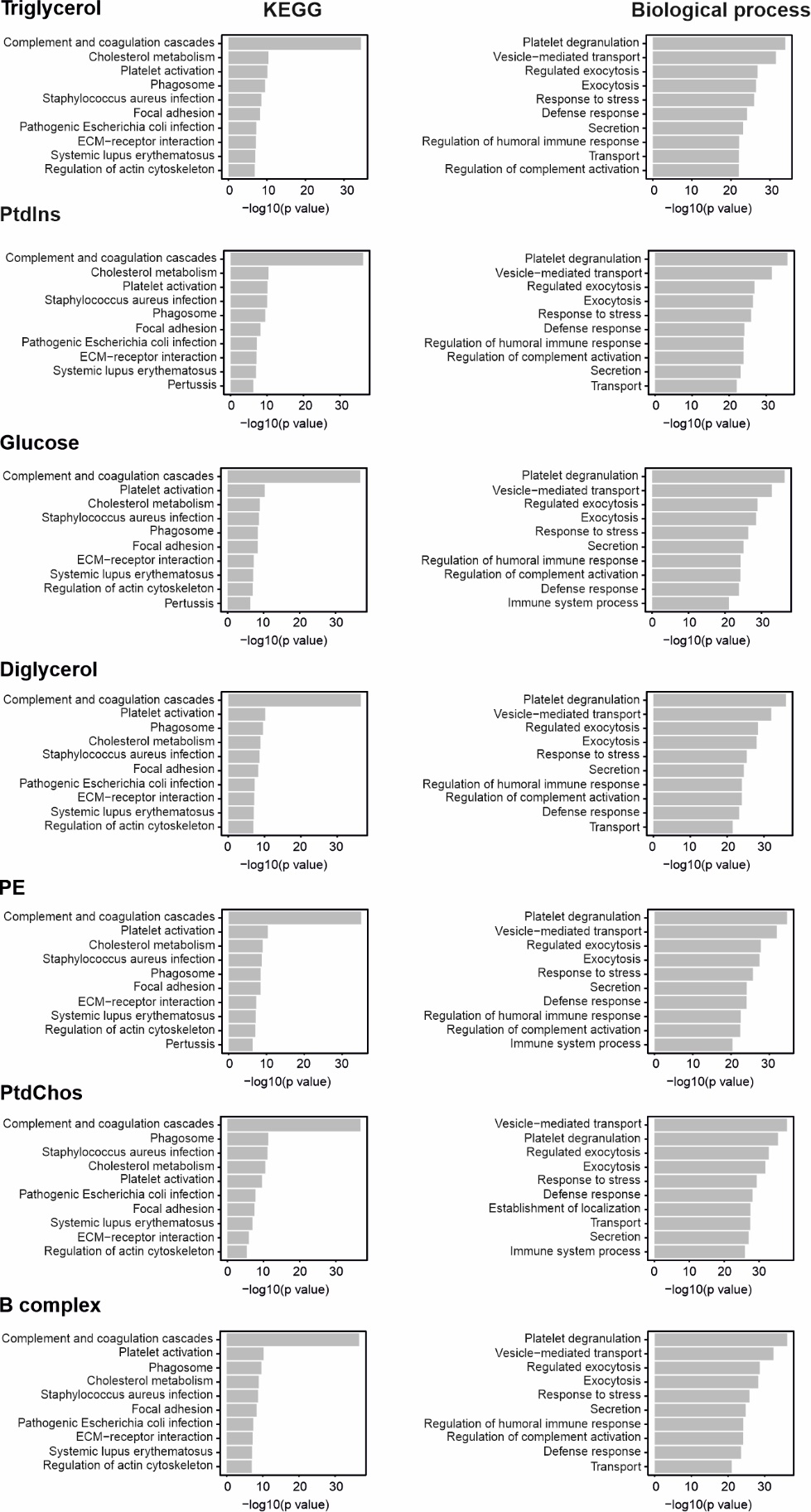

**Supplementary Fig. 5. Pathway enrichment for all significantly enriched and depleted proteins for each small molecule cumulatively across all concentrations.** KEGG and biological processes are shown.

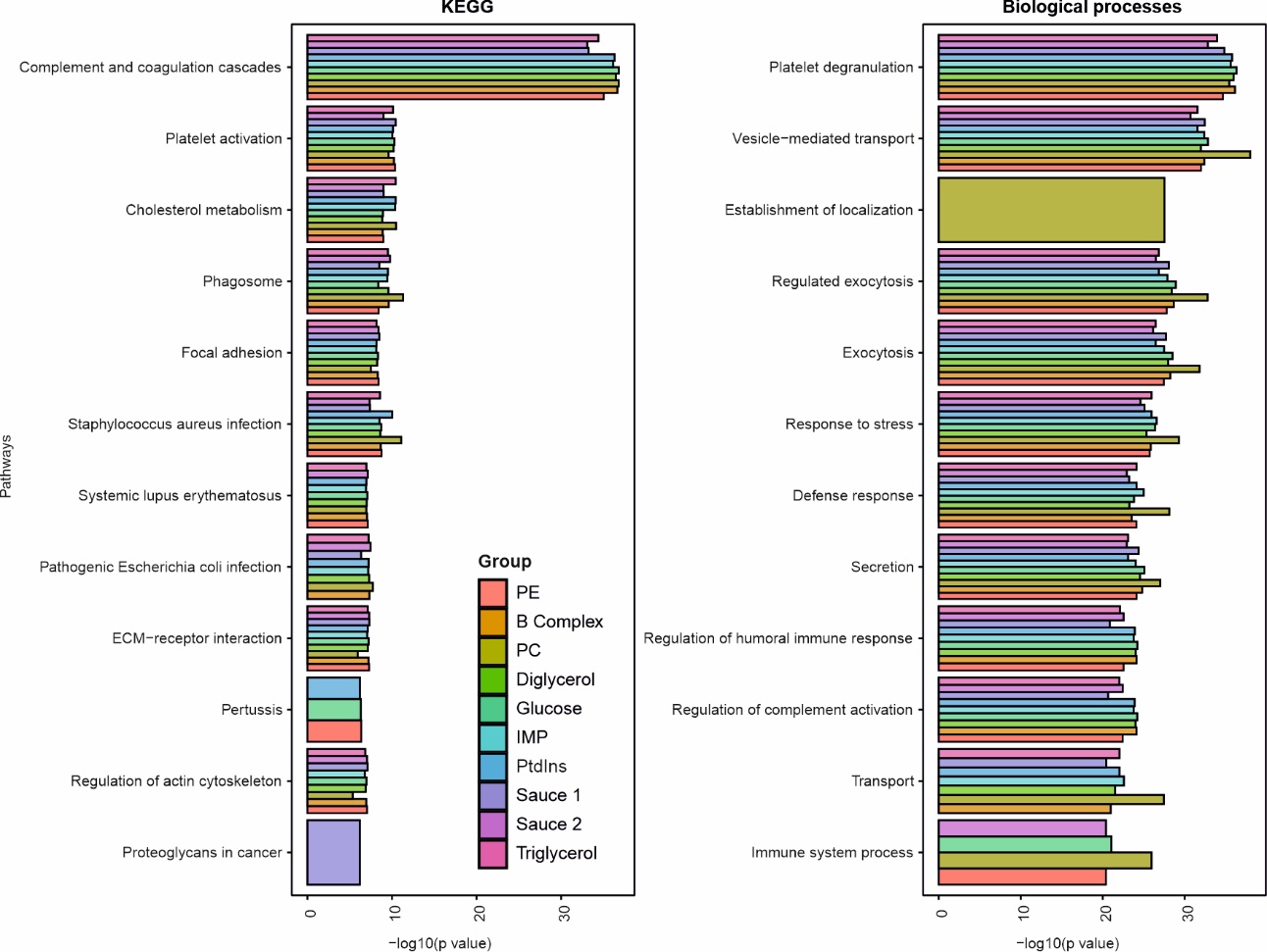

**Supplementary Fig. 6.** The combined enrichment plot for all small molecules and molecular sauces vs. untreated protein corona, cumulatively across all concentrations.

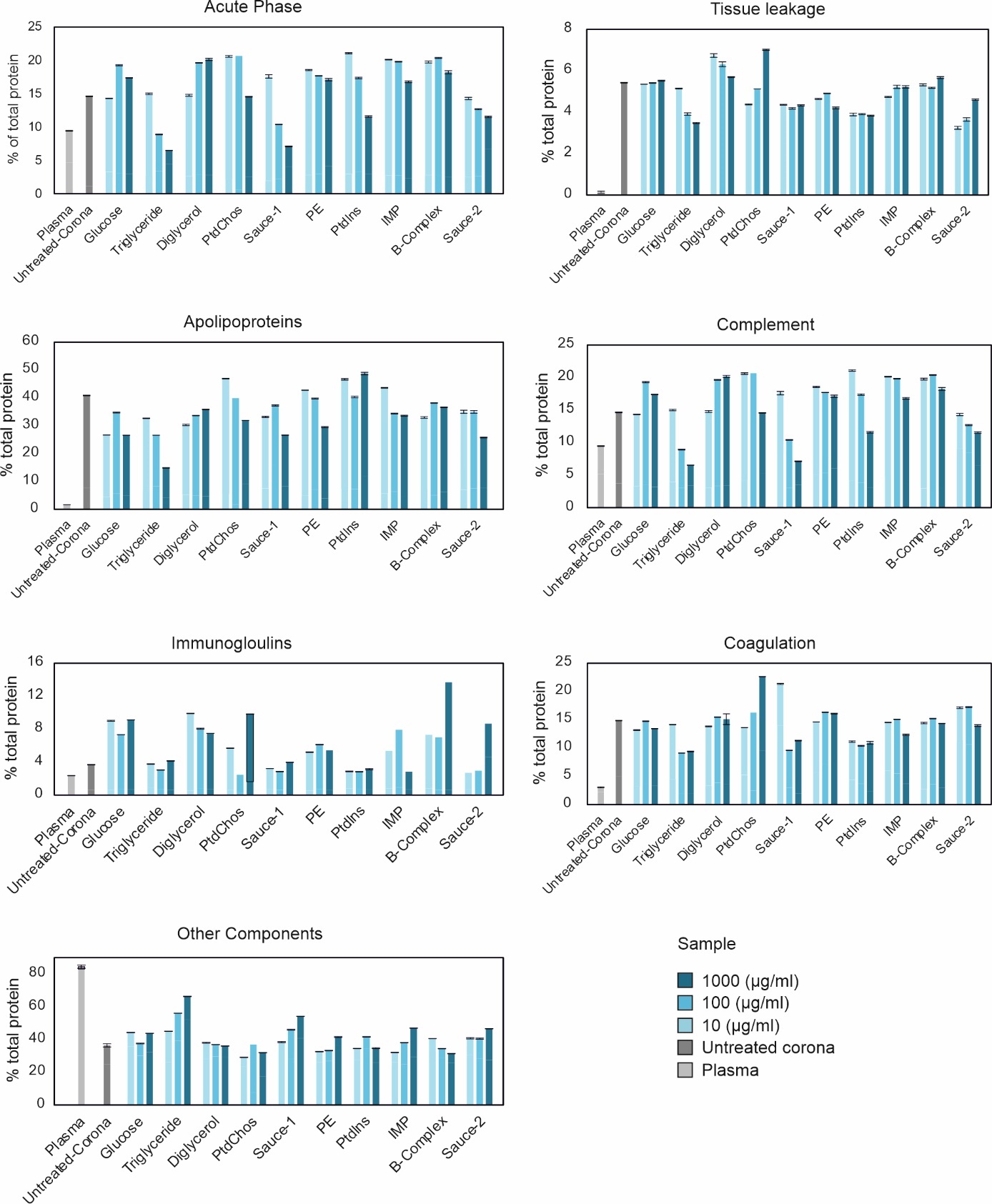

**Supplementary Fig. 7. Classification of quantified protein corona of various small molecules according to their physiological functions.**

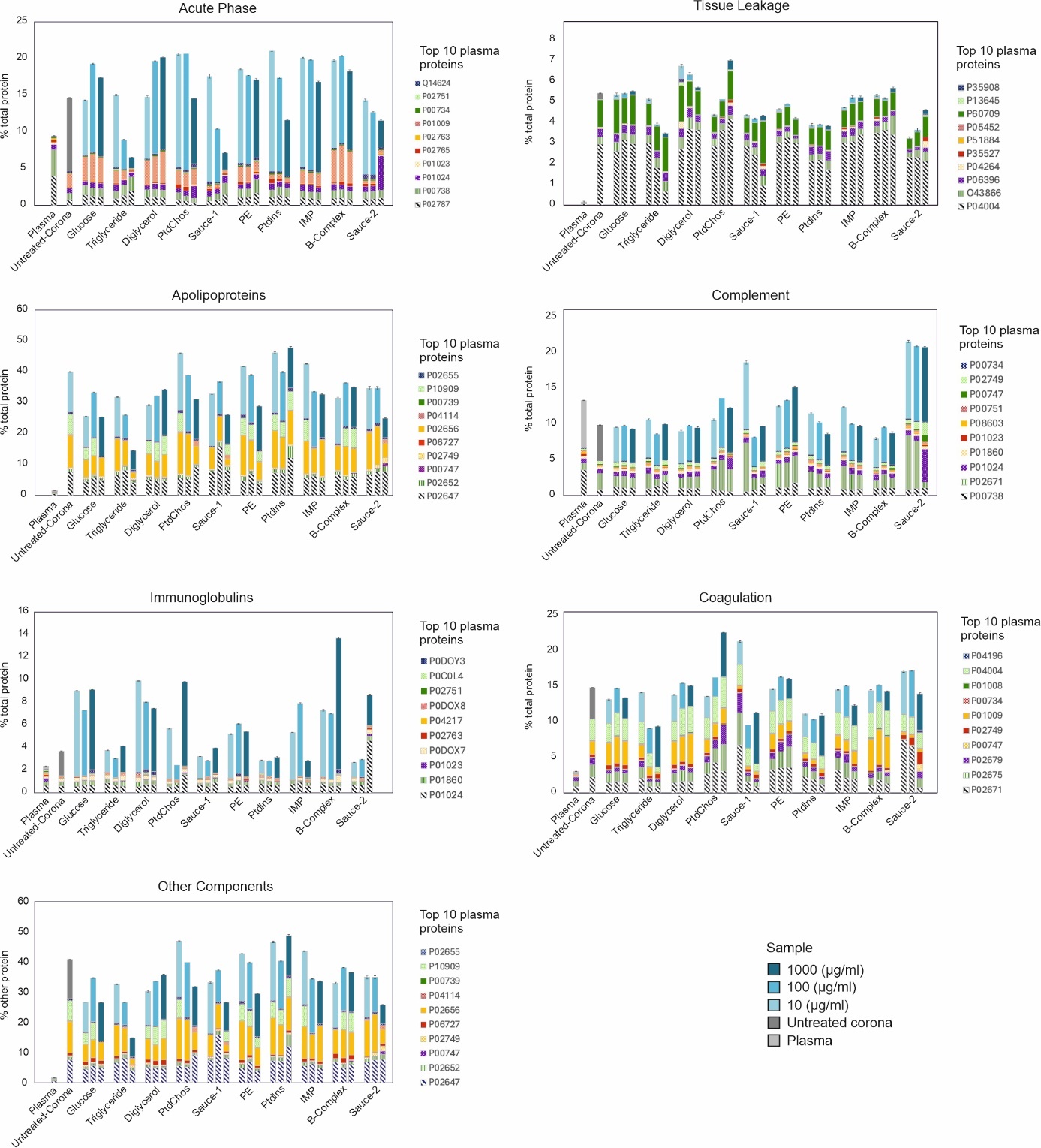

**Supplementary Fig. 8. Variations of top 10 plasma proteins across protein corona compositions of various small molecules according to their physiological functions.**

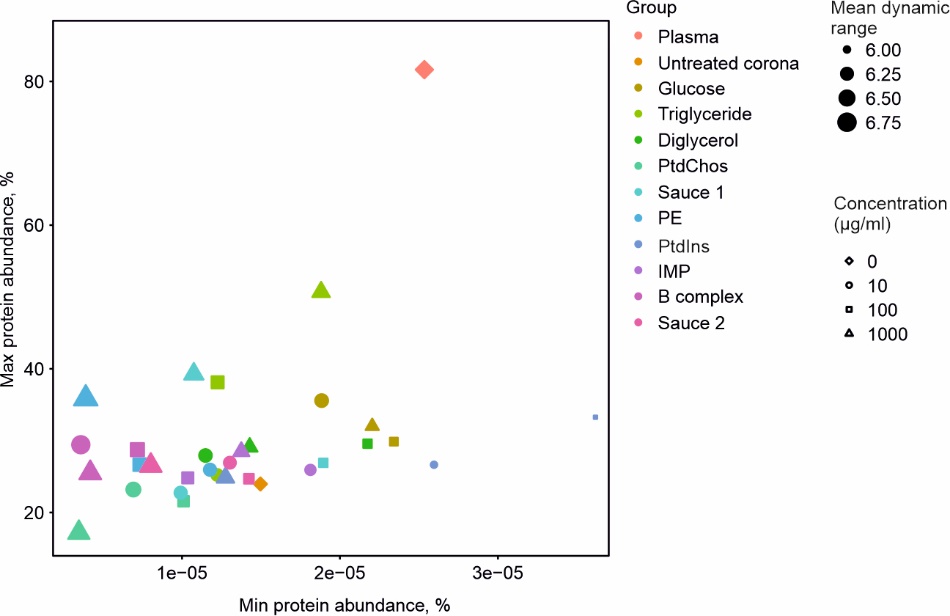

**Supplementary Fig. 9. The impact of spiking small molecules on the proteome dynamic range.** The dynamic range (order of magnitude) of the proteomics analysis for different samples is shown.

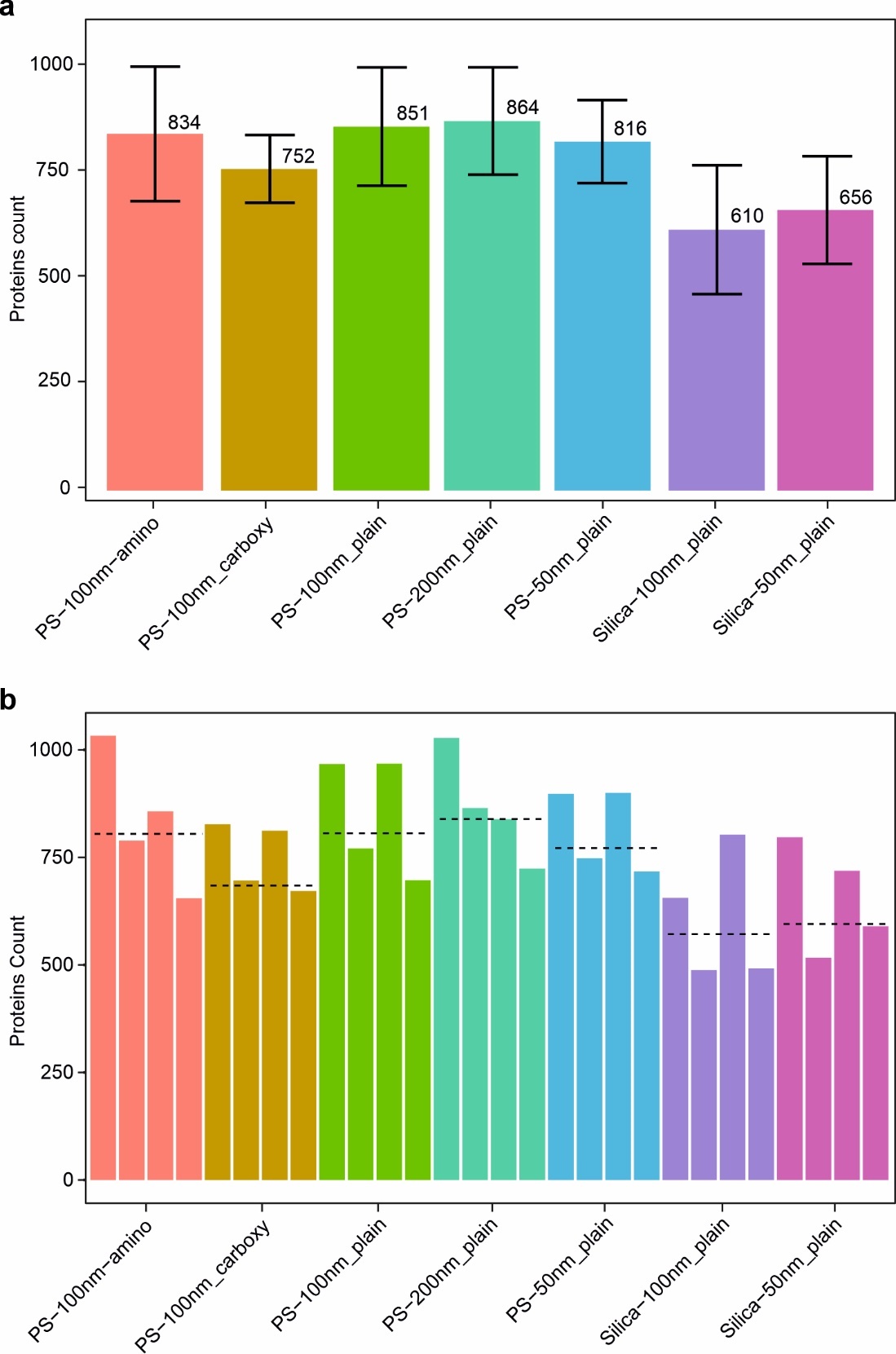

**Supplementary Fig. 10.** The impact of NPs of different composition, size and charge. **a,** The association of using various NPs with different sizes and charges with average protein count using plasma from 4 individual donors. Error bars represent mean±SD. **b,** same as panel a, but protein count shown for the 4 individual donors separately for each NP type. The line represents the average protein count.

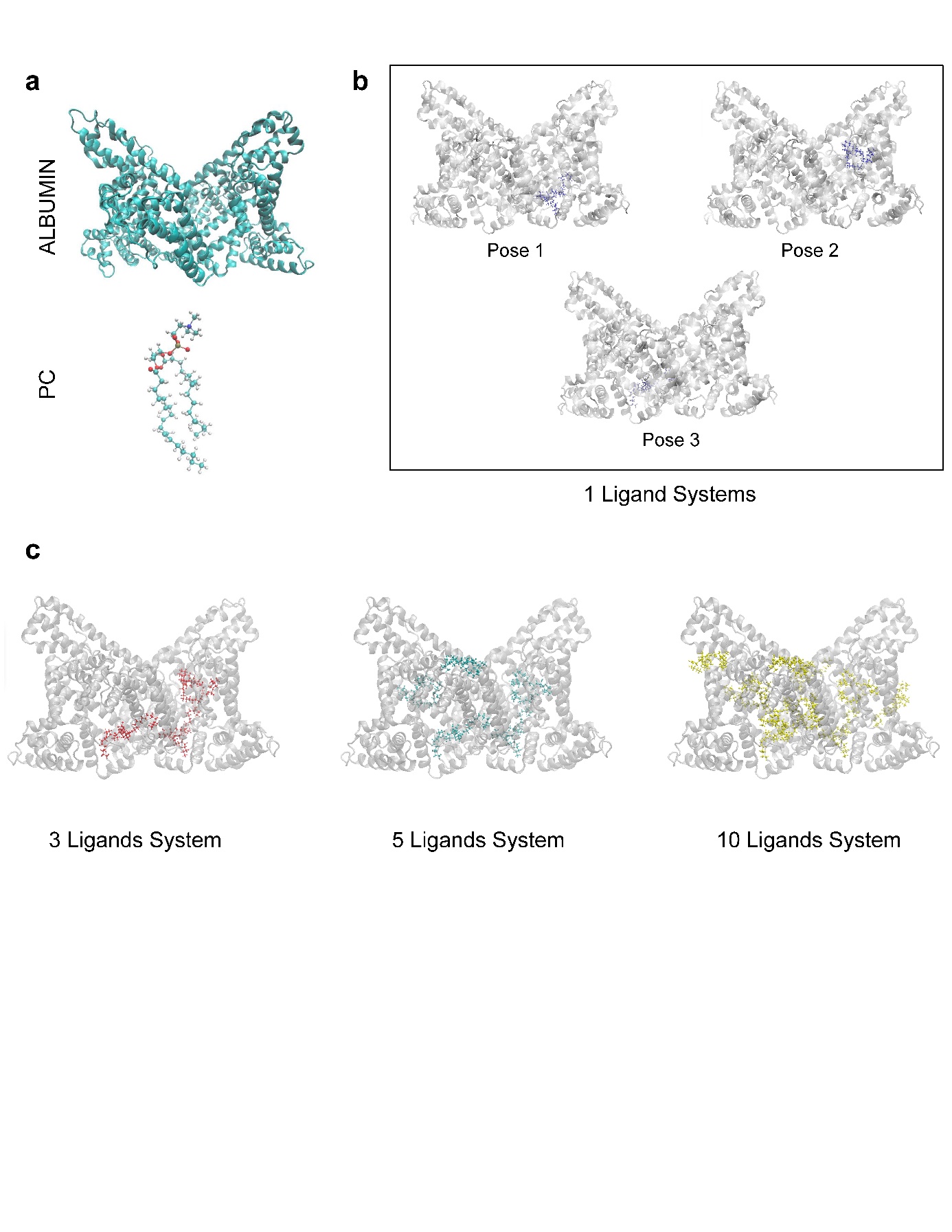

**Supplementary Fig. 11.**  **All-atoms molecular dynamics simulations components. a,** Crystal structure of albumin and representation of a single PtdChos (PC) molecule. **b,** Crystal structure of albumin showcasing the 1 ligand systems composed of the top 3 favorable binding sites for PtdChos as determined by molecular docking. **c,** Crystal structure of albumin illustrating the 3, 5, and 10 ligands systems which represent the the top 3, 5, and 10 most favorable binding sites for PtdChos respectively.

**Supplementary Tables**

**Supplementary Table 1.** The number of quantified proteins as well as the SD and CV of the number of quantified proteins for plasma, untreated protein corona and protein corona treated with various small molecules and molecular sauces (3 technical replicates)

| **Group** | **Concentration** | **Protein count** | **SD** | **CV (%)** |
| --- | --- | --- | --- | --- |
| Plasma | NA | 218 | 0.57735 | 0.265245 |
| Untreated corona | NA | 681 | 2.645751 | 0.38851 |
| Glucose | 10 | 615 | 6.110101 | 0.994051 |
| Glucose | 100 | 583 | 2.516611 | 0.431913 |
| Glucose | 1000 | 568 | 5.686241 | 1.001687 |
| Triglyceride | 10 | 682 | 3.605551 | 0.528673 |
| Triglyceride | 100 | 622 | 3.605551 | 0.579671 |
| Triglyceride | 1000 | 632 | 4.358899 | 0.689699 |
| Diglycerol | 10 | 665 | 2.309401 | 0.347104 |
| Diglycerol | 100 | 596 | 2.309401 | 0.387267 |
| Diglycerol | 1000 | 572 | 4.358899 | 0.762045 |
| PtdChos | 10 | 607 | 5.859465 | 0.965846 |
| PtdChos | 100 | 474 | NA | NA |
| PtdChos | 1000 | 897 | 3 | 0.334448 |
| Sauce 1 | 10 | 510 | 1.527525 | 0.299711 |
| Sauce 1 | 100 | 473 | 1.732051 | 0.366184 |
| Sauce 1 | 1000 | 622 | 1.527525 | 0.245451 |
| PE | 10 | 576 | 1 | 0.173611 |
| PE | 100 | 603 | 2.645751 | 0.438765 |
| PE | 1000 | 578 | 1 | 0.17301 |
| PtdIns | 10 | 434 | 3 | 0.691244 |
| PtdIns | 100 | 397 | 2.516611 | 0.63444 |
| PtdIns | 1000 | 410 | 5.131601 | 1.252629 |
| IMP | 10 | 550 | 2.081666 | 0.378714 |
| IMP | 100 | 717 | 1.527525 | 0.212945 |
| IMP | 1000 | 538 | 4.932883 | 0.917461 |
| B complex | 10 | 613 | 3.21455 | 0.524111 |
| B complex | 100 | 533 | 5.686241 | 1.067505 |
| B complex | 1000 | 552 | 4 | 0.724638 |
| Sauce 2 | 10 | 459 | 7.023769 | 1.531345 |
| Sauce 2 | 100 | 567 | 5.507571 | 0.971924 |
| Sauce 2 | 1000 | 558 | 2.645751 | 0.474149 |

**Supplementary Table 2.** The number of quantified proteins as well as the SD and CV of the number of quantified proteins for plasma, untreated protein corona and protein corona treated with various concentrations of PtdChos (3 technical replicates)

| **Group** | **Concentration** | **Average protein count** | **SD** | **CV (%)** |
| --- | --- | --- | --- | --- |
| Choline | 100 | 818 | 1.154701 | 0.141104 |
| Choline | 500 | 836 | 2.516611 | 0.30091 |
| Choline | 1000 | 957 | 3.05505 | 0.319121 |
| Choline | 10000 | 856 | 3.05505 | 0.356759 |
| Untreated corona | NA | 617 | 12.12436 | 1.96505 |
| Plasma | NA | 294 | 1.732051 | 0.589133 |

**Supplementary Table 3.** The number of quantified proteins as well as the SD and CV of the number of quantified proteins for protein corona treated with PtdChos and incubated with different NPs (plasma from 4 different donors)

| **Sample** | **Protein count** | **Average protein count** | **SD** | **CV (%)** |
| --- | --- | --- | --- | --- |
| PS_100nm_Amino_PtdChos_donor1 | 1033 | 834 | 157.263 | 18.86779 |
| PS_100nm_Amino_PtdChos_donor2 | 789 | 834 | 157.263 | 18.86779 |
| PS_100nm_Amino_PtdChos_donor3 | 857 | 834 | 157.263 | 18.86779 |
| PS_100nm_Amino_PtdChos_donor4 | 655 | 834 | 157.263 | 18.86779 |
| PS_100nm_Carboxylated_PtdChos_donor1 | 827 | 752 | 79.0796 | 10.5194 |
| PS_100nm_Carboxylated_PtdChos_donor2 | 696 | 752 | 79.0796 | 10.5194 |
| PS_100nm_Carboxylated_PtdChos_donor3 | 812 | 752 | 79.0796 | 10.5194 |
| PS_100nm_Carboxylated_PtdChos_donor4 | 672 | 752 | 79.0796 | 10.5194 |
| PS_100nm_PtdChos_donor1 | 967 | 851 | 138.1554 | 16.23925 |
| PS_100nm_PtdChos_donor2 | 771 | 851 | 138.1554 | 16.23925 |
| PS_100nm_PtdChos_donor3 | 968 | 851 | 138.1554 | 16.23925 |
| PS_100nm_PtdChos_donor4 | 697 | 851 | 138.1554 | 16.23925 |
| PS_200nm_PtdChos_donor1 | 1028 | 864 | 125.3289 | 14.50566 |
| PS_200nm_PtdChos_donor2 | 865 | 864 | 125.3289 | 14.50566 |
| PS_200nm_PtdChos_donor3 | 839 | 864 | 125.3289 | 14.50566 |
| PS_200nm_PtdChos_donor4 | 724 | 864 | 125.3289 | 14.50566 |
| PS_50nm_PtdChos_donor1 | 898 | 816 | 96.96176 | 11.88621 |
| PS_50nm_PtdChos_donor2 | 748 | 816 | 96.96176 | 11.88621 |
| PS_50nm_PtdChos_donor3 | 900 | 816 | 96.96176 | 11.88621 |
| PS_50nm_PtdChos_donor4 | 717 | 816 | 96.96176 | 11.88621 |
| Si_100nm_PtdChos_donor1 | 656 | 610 | 150.7456 | 24.72253 |
| Si_100nm_PtdChos_donor2 | 488 | 610 | 150.7456 | 24.72253 |
| Si_100nm_PtdChos_donor3 | 803 | 610 | 150.7456 | 24.72253 |
| Si_100nm_PtdChos_donor4 | 492 | 610 | 150.7456 | 24.72253 |
| Si_50nm_PtdChos_donor1 | 797 | 656 | 125.866 | 19.19421 |
| Si_50nm_PtdChos_donor2 | 517 | 656 | 125.866 | 19.19421 |
| Si_50nm_PtdChos_donor3 | 719 | 656 | 125.866 | 19.19421 |
| Si_50nm_PtdChos_donor4 | 590 | 656 | 125.866 | 19.19421 |
